# Star-Motifs: Revealing Single-Cell Spatiotypes from Routine Histology Using Star-Convex Neighborhoods

**DOI:** 10.1101/2025.11.07.687248

**Authors:** Gozde N. Gunesli, Muhammad Dawood, Lawrence S. Young, Fayyaz Minhas, Shan E Ahmed Raza, Nasir M. Rajpoot

## Abstract

The tumor microenvironment (TME) is a dynamic interplay among cancer, immune, and stromal cells that profoundly influences tumor growth, progression, and treatment response. Although recent spatial transcriptomics and multiplex imaging technologies offer fine-grained insights into the TME, their high cost and lengthy turnaround times limit routine clinical use. In contrast, whole slide images (WSIs) of Hematoxylin and eosin (H&E)-stained are readily available for most patients and provide valuable information about tumor biology. Here, we present a method that explicitly models each cell’s local neighborhood to identify recurrent spatial patterns, which we call “Star-Motifs.” We first encode the local microenvironment of each cell using a “Star-Environment” representation that captures distances and angles to multiple surrounding cell types. We then perform unsupervised clustering of these descriptors to discover distinct Star-Motifs. We validated our method on 1.8 billion cells across 3,105 diagnostic slides from The Cancer Genome Atlas, spanning five distinct cell types: neoplastic, inflammatory, stromal, necrotic, and non-neoplastic. Our method uncovered a diverse range of Star-Motifs, from densely packed tumor clusters to tumor cells surrounded by immune cells (tumor-infiltrating lymphocytes). At the patient level, the relative abundances of these Star-Motifs correlate with patient survival outcomes, immune subtypes, and molecular alterations, offering interpretable, clinically relevant insights. Moreover, classical machine learning models trained on these abundances match or surpass deep learning approaches in predicting overall survival and immune-related biomarkers while remaining interpretable. By capturing higher-order spatial arrangements from routinely available histology slides, Star-Motifs provides a scalable, transparent platform for TME profiling, biomarker discovery, and personalized risk assessment in oncology.

## 1 Introduction

The tumor microenvironment (TME) is widely recognized as a critical driver of tumor growth, progression, and therapy response, encompassing diverse immune and stromal cells, blood vessels, secreted factors, and the extracellular matrix [1–3]. With cells being the fundamental building blocks of tissues and organs [4] and fundamental components of TME, understanding their spatial organization which involves how they are arranged, distributed, and interact within the tissue, and understanding how the different regionally heterogeneous characteristics of these spatial organizations are associated with the outcomes, may provide key insights into cancer behavior and inform therapeutic strategies.

Earlier studies relied on bulk genomic and transcriptomic analyses to identify TME subtypes with distinct molecular and immune responses [5]. However, these methods lack the spatial context necessary to capture the full complexity of tumor dynamics [6]. Recent advances in spatially resolved omics technologies, such as multiplexed tissue imaging, enable high-resolution characterization of the cells and their interactions within the spatial context of the tumor, providing fine grained cellular insights into disease subtypes, therapeutic responses, and clinical outcomes [6–13]. However, these methods still face practical constraints related to cost, throughput, and seamless integration into clinical workflows [10]. Additionally, these technologies though provide fine grained cellular information they still need methods to identify spatial cellular arrangements driving the disease or that could stratify patients based on these spatiotypes (spatial cellular arrangements) [14].

In routine clinical practice, Hematoxylin & eosin stained (H&E) whole slide images (WSIs) remain the gold standard for cancer diagnosis and offer the advantage of large-scale data repositories such as The Cancer Genome Atlas (TCGA) [15] with various clinical and genomic ground truth labels. These histology images contain abundant information, and deep learning (DL) methods, namely multiple-instance learning (MIL) approaches and graph neural networks (GNN), have shown that H&E-based features can predict patient survival [16], various molecular and genetic alterations [17–19], drug sensitivities [20], and transcriptomic states [21]. From these approaches, while MIL methods ignore spatial relationships, GNNs are claimed to capture cellular topology. However, when capturing long-range spatial relationship GNNs could be prone to oversmoothing and subsequently losing interpretability [22]. In general, DL approaches operate as “black boxes” providing limited interpretability and insight into the underlying biological mechanisms [23–25].

Beyond DL methods, using histology images, numerous approaches [12, 26–32] have focused on spatial analyses of specific components of TME, especially the immune cells in the context of tumor-infiltrating lymphocytes (TILs) yielding clinically relevant biomarkers such as the ‘Immunoscore’ [26] and ‘sTIL’ [29–31]. These metrics typically assess TIL counts or densities within specific tumor or stromal compartments, offering prognostic value in multiple cancer types. However, they often rely on pre-selected regions, one or two cell-type categories and simple measures of spatial distribution [12], thereby capturing only a fraction of the TME’s complexity. Network-based approaches, in which cells are modeled as graph nodes and their spatial interactions as edges, can potentially reveal more complex intercellular relationships. However, these methods are sensitive to the chosen connectivity criteria (e.g., distance thresholds), and identification of higher-order motifs (recurrent patterns of multicellular architecture) from networks remains computationally infeasible [33]. While certain topological features (e.g., clustering coefficient, harmonic centrality) can indeed be extracted from cell graphs [34, 35], the biological interpretation of such features is not always straightforward.

In this study, we introduce a data-driven framework to uncover complex spatial organization of multiple cell types surrounding individual cells within the TME. Central to our approach is a novel representation of cell microenvironment (*Star-Environment*), which captures the spatial location and type information of surrounding cells around a cell of focus. Unsupervised clustering of these *Star-Environment* vectors yields *Star-Motifs*: recurrent spatial patterns that can be quantified and related to clinical and genomic features. Applying our *Star-Motifs Framework* to 3,105 diagnostic H&E WSIs spanning five TCGA cancer types, we identified 20 tumor associated motifs (NEO Star-Motifs) and 18 immune associated motifs (INF Star-Motifs). These motifs encompass all local micro-environmental arrangements, ranging from dense inflammatory infiltration to primarily neoplastic or stromal cell neighborhoods. For each patient we then compute patient-level relative motifs abundance and use it for downstream analysis. As shown in the following sections, Star-Motif frequencies reflect important biological variations within the TME, correlating with well-established image and transcriptomic based TME features such as TIL densities and immune expression signatures, several of them are prognostically relevant and can be used to stratify patients by survival outcome. Moreover, classical machine learning models trained on Star-Motif frequency vectors match or exceed the performance of deep learning methods on tasks such as survival prediction, predicting immune-related signatures, and specific genetic alterations, while providing an interpretable lens into tumor biology. To promote transparency and facilitate further research, we will publicly release the nuclear segmentation dataset generated in this study, comprising the locations, contours and cell-type labels of approximately 1.8 billion cells across all WSIs.

## 2 Results

### 2.1 Pan-cancer dataset and the Star-Motifs framework

In this work, we propose the *Star-Motifs Framework* that enables data-driven unsupervised discovery and characterization of higher-order spatial organization patterns on single-cell level. To extract these patterns and explore their clinical relevance and predictive utility, we applied this framework to a pan-cancer dataset collected from The Cancer Genome Atlas (TCGA) comprising 3105 H&E stained diagnostic wholeslide images (WSIs) from 2892 patients. Specifically, 1095 breast invasive carcinoma (BRCA), 585 colon and rectum adenocarcinoma (CRC), 511 lung adenocarcinoma (LUAD), 504 lung squamous cell carcinoma (LUSC) and 410 prostate adenocarcinoma (PRAD) WSIs were used. Each WSI underwent automated nuclei segmentation and classification using Hover-Net [36] model, which assigns individual nuclei to one of five cell types: Neoplastic epithelium (tumor), Inflammatory, Connective, Dead (Necrotic), and Non-neoplastic epithelium (normal)(Fig. 1A). This nuclei segmentation and classification step generated a large dataset comprising locations and types of approximately 1.8 billion detected cells (1,833,975,602 cells) out of which 43.2% were neoplastic epithelium, 10.4% were inflammatory, 21.6% were connective, 22.8% were dead, and 1.9% were non-neoplastic epithelium (Fig. 1B). This resulting dataset enabled in-depth quantitative analyses at single-cell resolution and formed the basis for subsequent steps focused on identifying higher-order spatial patterns within heterogeneous tumor tissues.

To explore pan-cancer spatial organization patterns, we developed the *Star-Environment* representation (Methods, Fig. 1C) which encodes the local neighborhoods of individual cells. Briefly, for a single cell’s local neighborhood, this representation is obtained by constructing multiple star-convex polygons, each representing the spatial arrangement of a different cell type surrounding that single cell at the center. For this, a polar coordinate system is centered on the cell of interest and divided into *N* equidistant angles (*N* = 360). For each cell type, the distance to the closest cell within a maximum radius is measured at each angle, resulting in a star-convex polygon contour that captures how that particular cell type is arranged around the center cell. We used a large profiling maximum radius value (500*µm* = 1000*pixels*), twice of the effective communication distance of a single cell (250*µm* [37]). Combining the star-convex polygons of all cell types yields a single vector, referred to as the *Star-Environment Vector*.

Finally, in a completely data-driven manner, we extracted different higher-order spatial organization patterns, which we call *Star-Motifs*, using the obtained *Star-Environment Vectors* (Fig. 1D). This is achieved by clustering the *Star-Environment Vectors* after low-rank approximation, and then aggregating rotation-variant clusters (Methods). We extracted the Star-Motifs separately for neoplastic and inflammatory cell environments and obtained two sets of motifs: NEO Star-Motifs and INF Star-Motifs. However, we should note that by design, the motifs around neoplastic cells (NEO Star-Motifs) already contain information related to all other cell types, including the inflammatory cells in the local microenvironment of neoplastic cells; similarly, the INF Star-Motifs contain information related to all other cell types including the neoplastic cells.

**Fig. 1.**
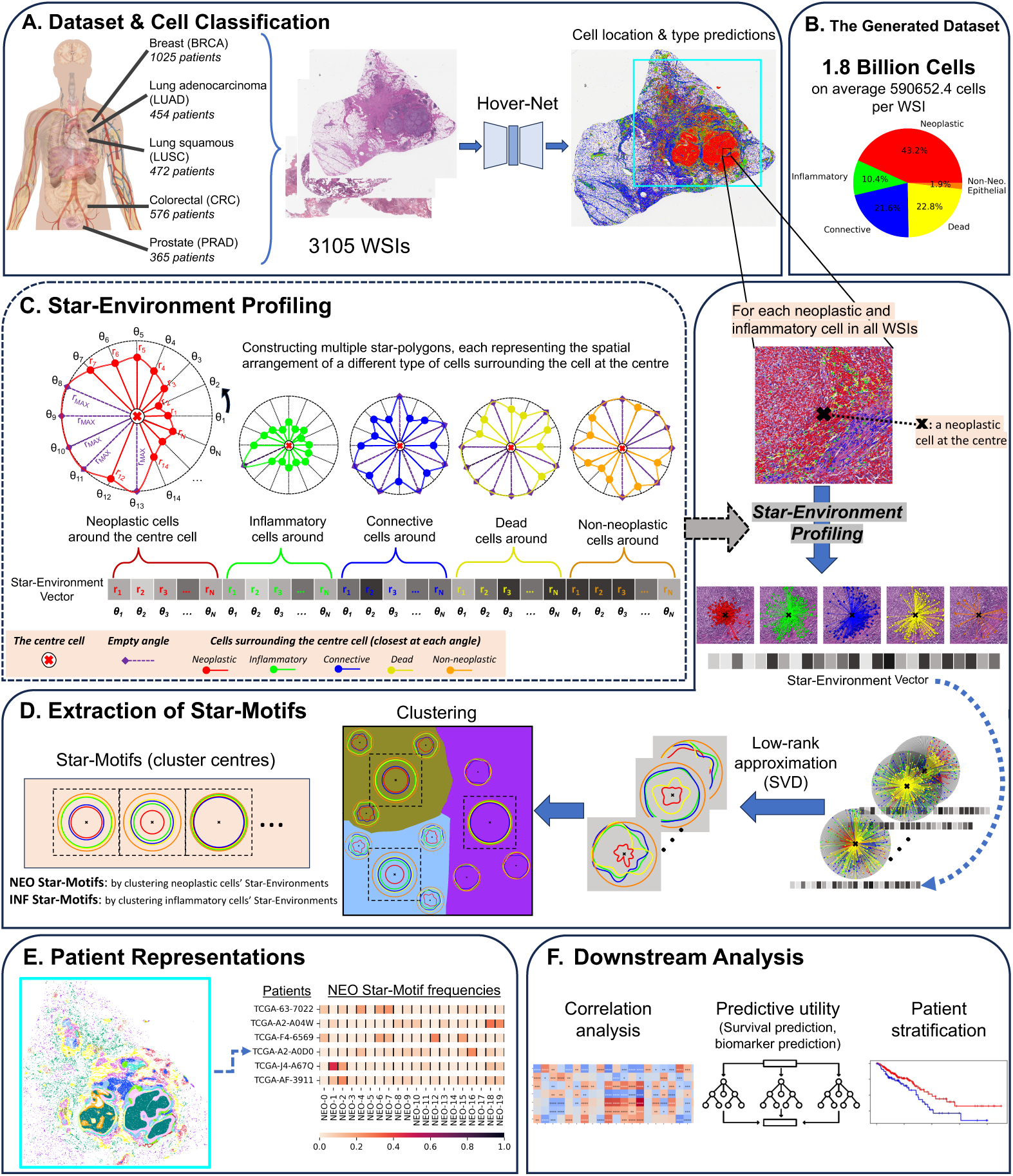
Overview of the *Star-Motifs* framework. **A** Whole Slide Images (WSIs) from the pancancer TCGA dataset are processed for nuclear segmentation and classification. **B** Cell composition of the entire dataset **C** Construction of a *Star-Environment* vector to model the spatial organization surrounding a cell at the center is demonstrated. **D** Schematic diagram of the data-driven extraction of *Star-Motifs*. NEO Star-Motifs are extracted by clustering *Star-Environments* of neoplastic cells, and INF Star-Motifs are extracted by clustering *Star-Environments* of inflammatory cells. **E** In each WSI, every neoplastic cell received a NEO Star-Motif label, and every inflammatory cell received an INF Star-Motif label. Then, patients (one or more WSI per patient) are represented by compositional vectors (Star-Motif frequency vectors) which account for the the fraction of cells assigned to each Star-Motif. Image illustrates the assigned NEO Star-Motif labels in the area inside the blue square in A. **F** Downstream analysis leverages the patient representations for integrative analysis and prediction.

In our analyses, we represented each patient by Star-Motif frequency vectors (Fig.1E, Methods). Specifically, for each WSI, every neoplastic cell received a NEO Star-Motif label, and every inflammatory cell received an INF Star-Motif label. We then computed the fraction of cells assigned to each Star-Motif for each patient (including multiple WSIs, if available). This yields two compositional vectors per patient, one for neoplastic cells (twenty dimensional) and one for inflammatory cells (eighteen dimensional), where each entry indicates the fraction of cells assigned to a particular Star-Motif. Our experiments and analyses (Fig. 1F) leveraged these patient-level representations.

### 2.2 Star-Motifs capture higher-order spatial organization patterns across cancer types

Through clustering of the *Star-Environments*, we identified 20 NEO and 18 INF Star-Motifs across all tumor types. These Star-Motifs were extracted directly from routine H&E-stained images without any prior assumptions on which local microenvironments might matter biologically. Except the Star-Motifs NEO-19 and INF-17 which are comprised of multiple clusters (merged based on rotation invariance, Methods), each one of these NEO and INF Star-Motifs correspond to one cluster, collectively capturing a range of higher-order spatial organization patterns of different cell types surrounding individual neoplastic or inflammatory cells.

To facilitate interpretation, we first reconstructed each cluster’s “star-convex polygons” in the original 2D space as illustrated in Figs.2A and S1A. With each color-coded star-convex polygon line corresponding to a specific cell type, the appearance of the multiple star-convex polygon lines in the illustrations represent the approximations of different patterns of different cell types surrounding a cell in the center and provide a quick overview of the spatial organization of different cell types each motif represents. For example, if we check the illustration for the Star-Motif NEO-1, we see that the red line (representing neoplastic cells around the center cell) and the blue line (representing the connective cells around the center cell) are closer to the center, while other lines (green, yellow, and orange) and correspondingly other cell types (inflammatory, necrosis, non-neoplastic epithelial) are represented farther away, implying a local-microenvironment more enriched in neoplastic and connective cells. We then performed a quantitative analysis (“Concentric enrichment analysis”, Methods) (Fig. 2C; NEO Star-Motifs all cell types in Fig. S2, and INF Star-Motifs all cell types in Fig. S1B), where we systematically examined, at different distances, how often each cell type was present around a center cell (in a Star-Environment) in each motif versus its average occurrence across all Star-Environments and calculated log_2_(fold enrichment) scores to assess whether a certain cell type is relatively over-represented (i.e., abundant, with more enrichment) or under-represented (i.e., less abundant, with less enrichment) in a motif at a particular distance. This quantitative analysis not only corroborates the visual polygon-based observations but also illuminates subtle differences, whether enrichments of different cell types differ with distance around a given Star-Motif, such as how certain Star-Motifs feature inflammatory cells at short ranges(e.g., NEO-17, NEO-18) or at long ranges (e.g., NEO-1, NEO-7).

**Fig. 2.**
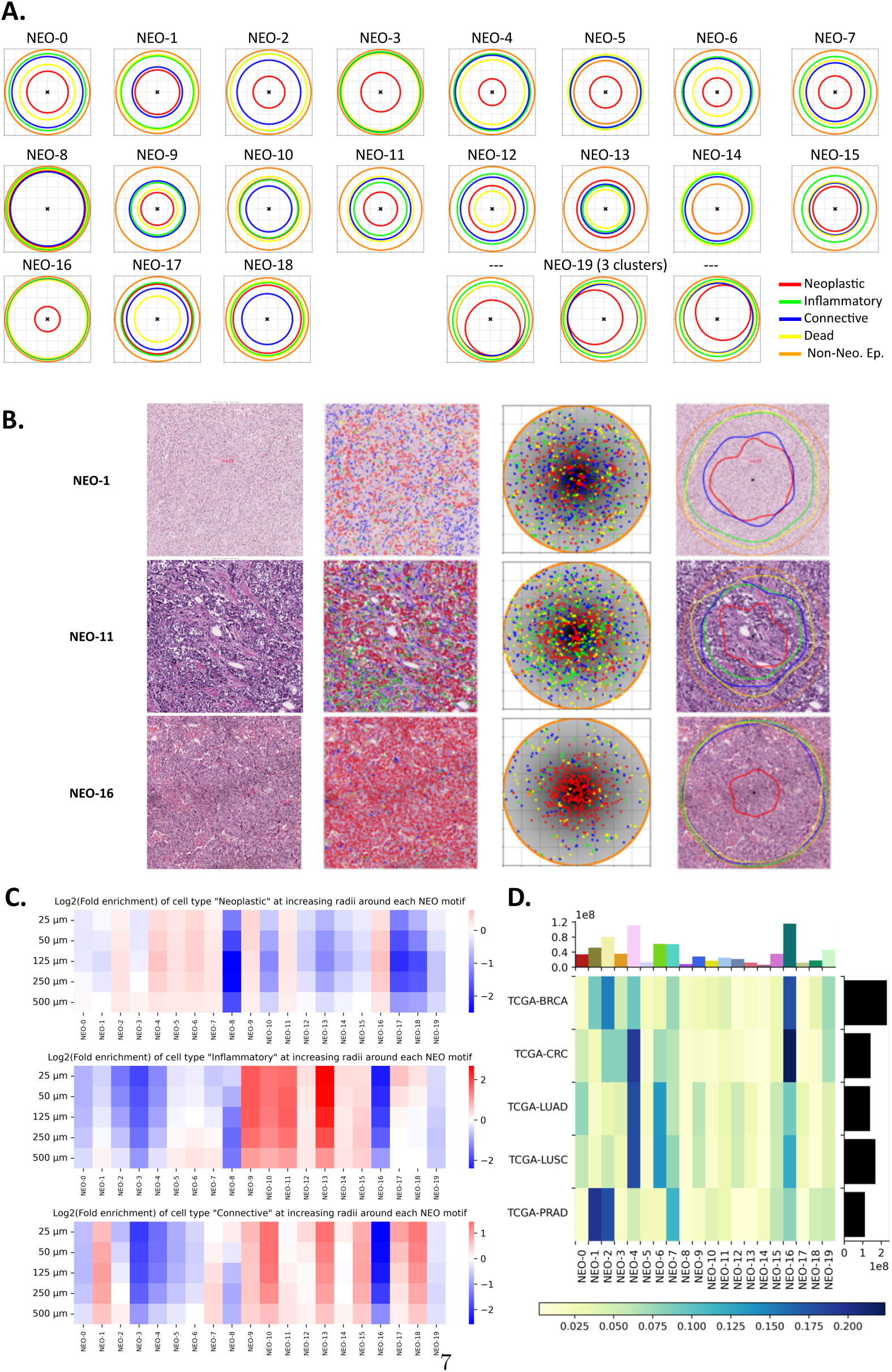
NEO Star-Motifs. **A** Extracted NEO Star-Motifs are illustrated. Each motif in the figure shows the reconstructed cluster center from the low-rank space to the original space. These Star-Motifs, which are obtained in a data-driven manner, represent the approximations of different higher-order patterns of different cell types surrounding a cell in the center (with the center cell being a neoplastic cell in NEO Star-Motifs). **B** Most representative Star-environments (closest to the cluster centers). Shows image patches with the cells having the most representative environment at the patch center (column 1), corresponding nuclei segmentation (column 2), Star-environments before (column 3) and after (column 4, overlaid on image patches) low-rank transformation. **C** The number of different types of cells surrounding each NEO cell with increasing concentric window sizes. Showing log(fold enrichment) values of cell types at each distance for each NEO motif. **D** NEO Star-Motifs VS cancer type. Heatmap showing the fraction of cells in each of the twenty NEO Star-Motifs in each cancer type. Rows were normalized to sum to 1. Histograms show the total number of cells exhibiting each motif and in each cancer type. All neoplastic cells were included to compute the heatmap and the histograms.

Overall, these Star-Motifs span a broad spectrum of higher-order relationships in the local microenvironments. Some motifs were predominantly enriched in tumor cells (NEO-3 and NEO-16; INF-14); whereas others (NEO-5, NEO-14, INF-16) show characteristically higher abundance of non-neoplastic epithelium. A few motifs exhibited low abundance of all cell types (NEO-8, INF-1), while some others (e.g., NEO-18, INF-7) were particularly enriched in connective cells. Inflammatory cells were overrepresented in multiple motifs (NEO-9,10,11,12,13,14,15; INF-2,6,8,9,10,11), but they differed in the extent of neoplastic, stromal, necrotic and/or non-neoplastic involvement within that inflammatory context. For example, NEO-9 and NEO-13, while they differ from each other in neoplastic cell enrichment, they also show more necrotic cells compared to NEO-11 or NEO-10. Among these high inflammatory Star-motifs, all but NEO-11 and INF-8 had over-represented connective cells in close proximity, indicating different variants of inflammatory and stromal co-enrichment captured by these motifs (NEO-9,10,12,13,14,15; INF-2,6,9,10,11).

To validate that the extracted Star-Motifs reflect spatial organization patterns in the tissue, we examined image patches cropped around cells whose Star-Environments are closest to each cluster’s centroid (Figs. 2B and S1C, and additional examples from each cancer type in Fig. S3.) and we observed that the cellular arrangements in actual tissue sections were consistent with motif interpretations.

We analyzed the fraction of cells exhibiting each Star-Motif in each cancer type (NEO Star-Motifs in Fig. 2D, INF Star-Motifs in Fig.S1D) and Star-Motif frequency distributions of patients in each cancer type (Fig.S4). While every Star-Motif existed in every cancer type, they were not distributed in an equal manner among cancer types, possibly indicating spatial organization characteristics varying by tumor type. For example, motifs dominated by neoplastic cells alone (NEO-3, NEO-16), indicating dense tumor regions, showed higher abundance in CRC, potentially reflecting glandular epithelial structures with fewer stromal or inflammatory neighbors.

We observed that certain Star-Motif frequencies in patients’ data were strongly correlated (Fig. S5), indicating that when one motif is abundant, another often is as well. In particular, strong cross-set correlations (i.e. between NEO and INF Star-Motifs) observed when both motifs captured similar spatial contexts. Because both NEO and INF Star-Motifs encode information on multiple cell types, including neoplastic and inflammatory cells, in their local environments, such overlapping content likely explains the strong cross-set correlations (e.g., INF-16 with NEO-5, both having high non-neoplastic cell enrichment; Spearman *ρ* = 0.9). We also detected notable intra-set correlations, suggesting that different yet related microenvironment patterns frequently coexist. Among the NEO motifs, NEO-5 and NEO-14, both enriched in non-neoplastic cells—showed the strongest within-set correlation (*ρ* = 0.8), while other results varied. For instance, NEO-18 was positively correlated with NEO-1, 8, 10 (*ρ* = 0.6–0.7) and negatively correlated with NEO-4, 6, 16 (*ρ* = −0.4–−0.5), indicating that certain motifs often appear together while others may be more mutually exclusive. Overall, these correlation patterns might be related to the complexity of tumor ecology, where multiple distinct local microenvironments can coexist or exclude one another within a single patient.

### 2.3 Star-Motif frequencies capture TIL, stromal, and leukocyte content, and proliferation

To investigate whether Star-Motifs capture key tumor microenvironment (TME) features, we correlated patient-level Star-Motif frequencies with four established metrics provided by Thorsson *et al.*[5](Methods): *tumor-infiltrating lymphocyte (TIL) fraction*, derived from the TIL maps produced using deep learning on H&E slides by Saltz *et al.*[27]; *Leukocyte fraction (LF)*, which estimates overall leukocyte content using DNA methylation data; *Stromal fraction (SF)*, reflecting non-tumor (stromal) cell content; *Proliferation*, representing tumor growth dynamics. As shown in Figs. 3A and S6, we observed numerous significant (Kendall’s *τ*) correlations between Star-Motif frequencies and these metrics across multiple tumor types, indicating that the local spatial arrangements captured by Star-motifs reflect these characteristics.

Unsurprisingly, Star-Motifs more enriched in inflammatory cells (NEO-6, NEO-9, NEO-10, NEO-11, NEO-12, NEO-13, NEO-15; and INF-2, INF-6, INF-8, INF-9, INF-10, INF-11) exhibited weak (Kendall’s *τ >* 0.10) to strong (Kendall’s *τ >* 0.45) positive correlations with TIL fractions across all examined cancers, indicating that these particular local environments likely represent “immune-infiltrated” regions (Fig. 3A). Conversely, Star-Motifs with low overall cell abundance (NEO-8, INF-1), those enriched almost exclusively in neoplastic cells (NEO-3, INF-14), boundary-associated Star-Motifs with low abundance other cell types (NEO-19, INF-17) and ones showing closer representation of neoplastic and connective cells (NEO-1, NEO-2; INF-12, INF-15) were negatively associated with TIL fractions, highlighting tumor regions where immune cell infiltration is limited. Correlations between Star-Motifs and Leukocyte fraction followed a similar trend. To visually confirm our statistical findings, we overlaid the NEO Star-Motifs positively correlated with TIL fractions on each cancer type, and found that cells labeled with these Star-Motifs coincide with the TIL regions on the TIL maps produced by Saltz *et al.*[27](Fig. 4), further validating that the Star-Motifs identified by our unsupervised framework dissect the tumor microenvironment into biologically meaningful patterns such as immune infiltration. Significant associations with proliferation were particularly more common in breast cancer (BRCA). Star-Motifs positively associated with proliferation signature tended to have higher enrichment of neoplastic cells. However, subtle differences in micro-environmental composition might be important. For instance, although NEO-1 exhibited a higher enrichment of neoplastic cells than NEO-13, it was negatively correlated with proliferation in BRCA and PRAD, whereas NEO-13 (comparatively richer in necrotic and inflammatory cells) showed a positive association. This might suggest that differences in inflammatory and necrotic cell composition in the local micro-environment may be linked to variations in proliferative behavior and potentially reflect tumor aggressiveness, supporting our motivation to extract higher-order spatial organization patterns.

**Fig. 3.**
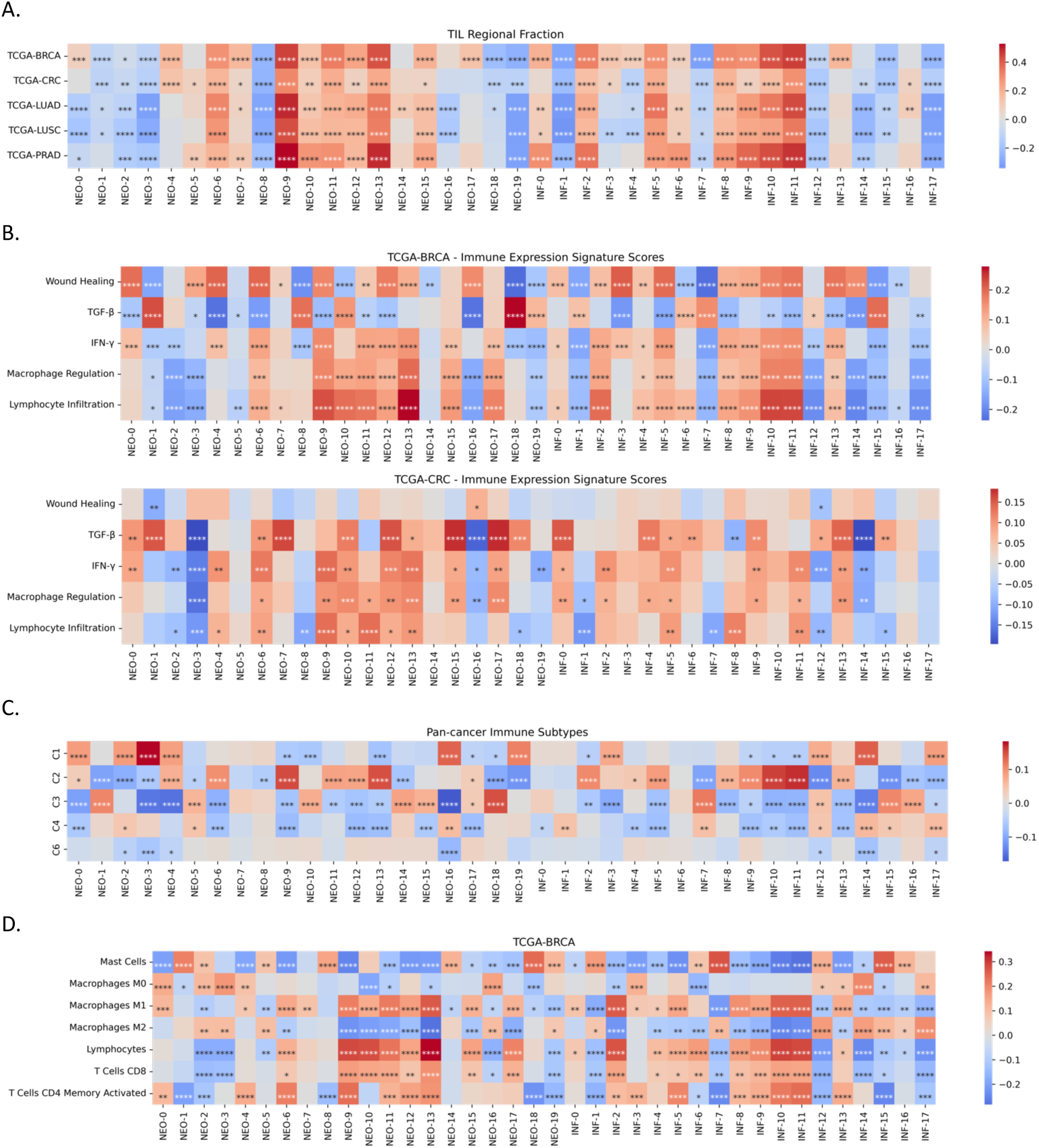
Star-Motif correlates of TME characteristics. **A.** Associations of motifs to TIL Regional Fractions across all cohorts. **B.** Associations between immune expression signature scores and motifs on BRCA and CRC cohorts **C.** Associations between motifs and pan-cancer immune subtypes C1, C2, C3, C4, and C6 (with number of patients 1122, 782, 569, 154, 79 respectively), **D.** Associations between immune cells and motifs on BRCA cohort. All heatmaps show Kendall’s *τ* correlation coefficients, BH correction for FDR. \**P ≤* 0.05, \*\**P ≤* 0.01, \*\*\**P ≤* 0.001, \*\*\*\**P ≤* 0.0001

### 2.4 Star-Motif frequencies reflect immune expression signatures and immune subtypes

We next investigated whether Star-Motif frequencies reflect transcriptomic immune expression signature scores (e.g., lymphocyte infiltration, macrophage regulation), which were used to cluster TCGA tumor types into immune subtypes (C1–C6) by Thorsson *et al.* [5]. Across all tumor types, we observed many significant associations with Immune Expression Signature Scores(Figs.3B and S7), although there were fewer associations in some cancer types (There were no significant associations with IFN *γ* and Wound healing in LUAD and, IFN *γ* in LUSC,). Correlations with immune response related signature scores, such as Macrophage Regulation, IFN *γ* and Lymphocyte Infiltration, tended to be concordant with associations to TIL regional fractions and positive associations were observed with inflammatory cell enriched Star-Motifs. Signatures including Wound healing and TGF-*β* demonstrated more complex or varied patterns.

**Fig. 4.**
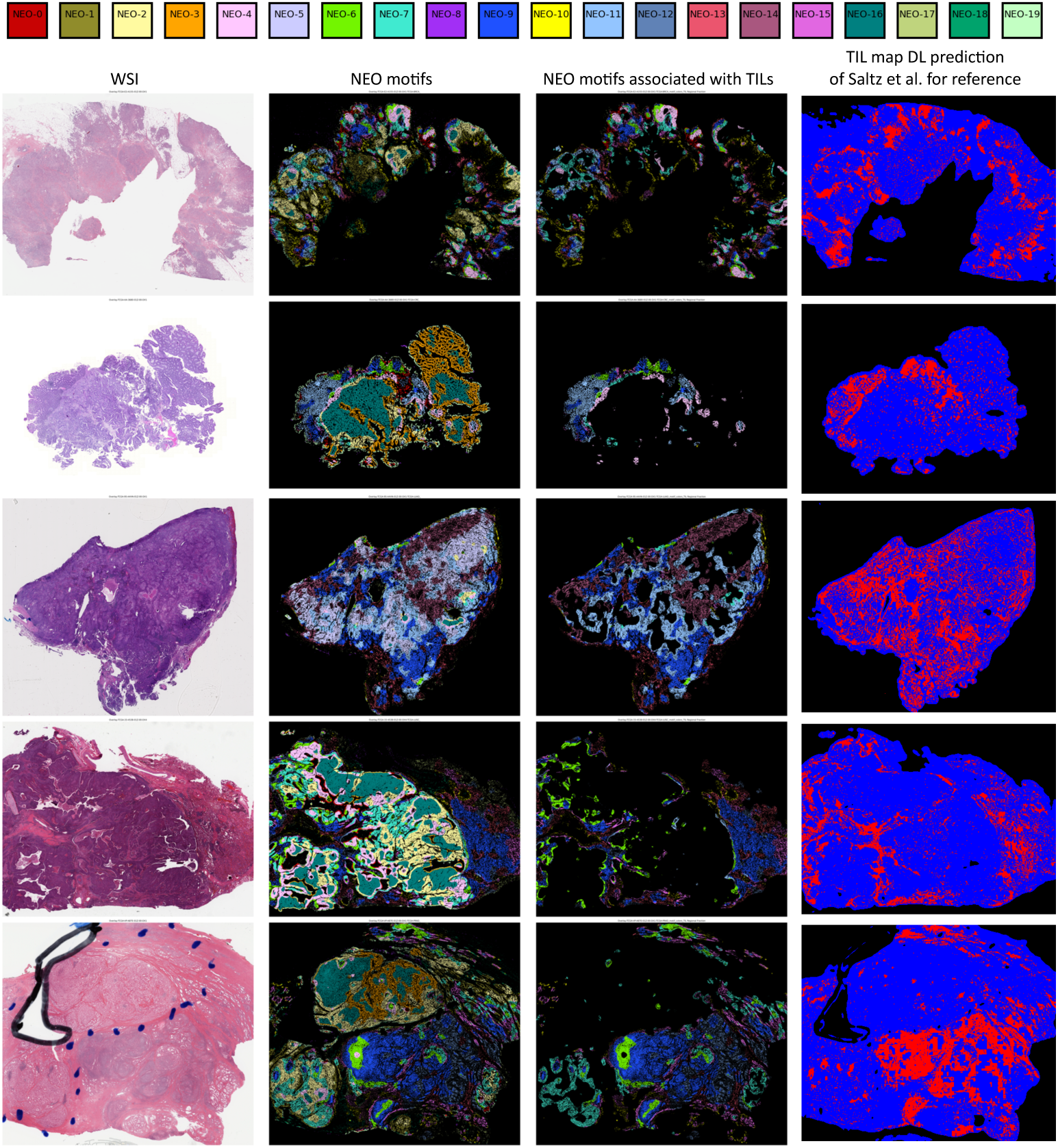
Example visual overlays of NEO Star-Motifs versus TIL regions in five WSIs (one per row) from BRCA, CRC, LUAD, LUSC, and PRAD. **Column 1:** Original H&E image. **Column 2:** Colorcoded map of all NEO Star-Motifs, with each neoplastic cell displayed in a unique color based on its assigned Star-Motif label, revealing the spatial distribution of different local microenvironments in neoplastic cells. **Column 3:** Same NEO Star-Motifs map, but restricted to the motifs that are significantly positively correlated with TIL fractions. **Column 4:** For reference, TIL maps produced using deep learning on H&E slides by Saltz *et al.*[27], where red indicates TIL-rich regions. These side-by-side comparisons show that Star-Motifs correlated with TIL regional fractions localize to the TIL regions. This further validates the ability of the Star-Motifs identified by our unsupervised framework to dissect the tumor microenvironment into biologically meaningful patterns such as immune infiltration.

**Fig. 5.**
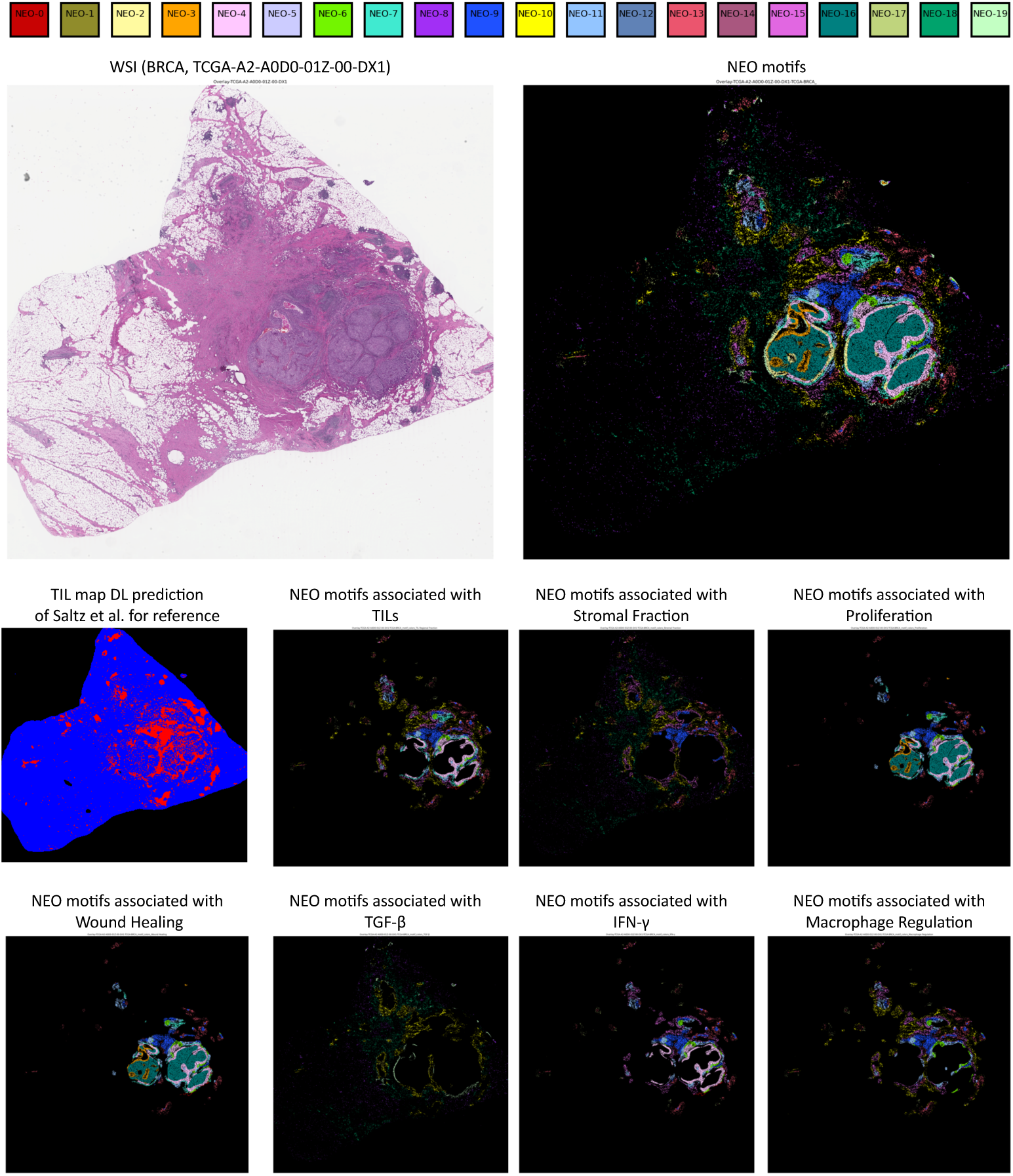
Visualization of Star-Motifs associated with different TME characteristics localize, on a single WSI (from TCGA-BRCA). Top row: (1) the original H&E image, and (2) a color-coded map of all NEO Star-Motifs, each neoplastic cell displayed in a color based on its assigned Star-Motif label. Subsequent rows: NEO Star-Motifs that are significantly correlated with eight different metrics, shown as separate overlays—namely, TIL Regional Fraction (with the Saltz *et al.*’s [27] TIL map prediction for reference), Stromal Fraction, Proliferation, Wound Healing, TGF-*β*, IFN-*γ*, and Macrophage Regulation. While transcriptomic-based TME metrics do not provide direct spatial information, these Star-Motif overlays provide a cell-level visualization showing which local microenvironments are linked to each biological process or immune function.

TGF-*β* signaling, which has a vital role in cancer by reshaping TME with immunosuppressive effects and demonstrated to be an impactful factor in poor response to immunotherapy [38], were mostly positively correlated with Star-Motifs enriched in connective cells (e.g., NEO-1,NEO-7,NEO-10,NEO-13,NEO-15,NEO-18). This observation is in line with the literature as TGF-*β* signaling is known to be associated with fibroblast proliferation and stromal fibrosis[38–40]. Also, previous studies which also extracted TME features from histology images [41, 42] demonstrated associations of TGF-*β* signaling with histology features related to high fibroblast density. Interestingly, some correlations differed by tumor type, showing tumor-type-specific correlations. For example: while both in BRCA and CRC, TGF-*β* signaling was significantly positively associated with NEO-1,10,18 which are very enriched in connective cells, certain motifs (NEO-0, NEO-6, NEO-12) exhibited opposite correlations between BRCA (negative) and CRC (positive), despite NEO-12’s association with stromal fraction (density) in both cancer types. So, the abundance of neoplastic cells having the same specific types of local microenvironment (the same motifs), had opposite relationships with TGF-*β* signaling in different cancer types. These observations suggest that a Star-Motif (i.e., a particular local microenvironment type) can exhibit contextdependent roles, while highlighting how motifs capture the heterogeneous and spatially organized nature of the TME in different cancer types.

Additionally, although these transcriptomic signatures lack direct spatial annotation, our framework pinpoints the local microenvironments linked to those signals. Therefore, we also visually examined how these motifs appear at the tissue level on a WSI (Fig. 5). We observed that Star-Motifs associated with TGF-*β* were localized to Stromal regions without immune infiltration.

Finally, we observed significant associations between Star-Motif frequencies and Thorsson *et al.*’s pan-cancer immune subtypes [5] (Fig. 3C). These immune subtypes integrate immune expression signature scores and correspond to distinct inflammatory or immunosuppressive states across tumor types. The most inflammatory subtype C2 (IFN-*γ* dominant) correlated with most of the inflammatory cell enriched motifs (NEO-9,11,12,13; INF-2,8,9,10,11). The wound-healing C1 subtype was positively associated with various motifs (NEO-0,2,3,4,16,19; INF-3,12,14,17). Notably, in all of these motifs inflammatory cells were under-represented. C3 subtype, which has the best prognosis over immune subtypes, were associated with few inflammatory cell enriched motifs (NEO-10,14,15), while we noticed that most motifs associated with this subtype had over-representation of non-neoplastic cells (NEO-1,5,10,14,15,18; INF-15,16). These correlations reinforce the idea that local spatial organization patterns captured and summarized by Star-Motif frequency vectors may reflect the nature of the immune responses in the corresponding tumor microenvironments.

### 2.5 Star-Motifs show associations with molecular estimates of different immune cells

We further examined whether there was evidence of variations in immune cell populations across Star-Motifs, including lymphocytes (overall, and also specifically for signatures of CD8 T cells and activated CD4 memory activated T cells), macrophage subsets, and mast cells. Although these cells play distinct roles in the tumor microenvironment, they cannot be directly distinguished in H&E images; instead, we used the estimates of their proportions which are available through analysis of the molecular data [5]. Associations of the Star-Motif frequencies with these molecular estimates of different immune cells provided insight into the functional difference of the local-microenvironments each Star-Motif represent (Figs.3D and S8). In general, Star-Motif frequencies showed varying patterns of association with T-cell (CD8+,CD4+ memory), lymphocyte, mast cell, and macrophage (M0, M1, M2) molecular estimates, showing that cells sharing the same morphological class (i.e., neoplastic or inflammatory) can exhibit different relationships with these immune cells depending on their specific local microenvironment (specific Star-Motif label).

In particular, many Star-Motifs that correlated positively with lymphocytes and specific T-cell proportions (e.g., NEO-9, NEO-10, NEO-11, NEO-12, NEO-13, NEO-15; INF-2, INF-8, INF-9, INF-10, INF-11) were mostly the ones with over-representation of inflammatory cells, whereas other associations (mast cells and macrophage subsets) seemed more complex. A closer look at tumor-associated macrophages (TAM) illustrates the heterogeneity in how motifs relate to M1, M2, and M0 subsets. While TAMs are known to differentiate according to their local microenvironment [43], earlier work on histology images has not reflected such heterogeneity (Discussion). In comparison, our Star-Motif approach revealed diverse associations, such that some motifs showed opposite associations with the pro-inflammatory M1 subtype and the tumor-promoting M2 subtype. While pro-inflammatory M1 subtype were positively associated with motifs with over-representation of inflammatory cells (e.g., NEO-9, NEO-10, NEO-11, NEO-12, NEO-13, NEO-15), these motifs had negative correlations with the M2 subtype. More diversity was observed with the undifferentiated M0 subtype. In CRC, for example, NEO-12 showed concurrent positive correlations with M0 and M1, whereas NEO-2 was positively associated with M0 yet negatively with M1. Moreover, the same TAM subset can display differing correlations across tumor types: M0 was positively associated with NEO-1 in CRC but negatively in BRCA, and NEO-16 exhibited the opposite pattern.

Thus, integrating molecular estimates of immune cell proportions with our Star-Motif Framework analysis suggested that immune cell subsets may have different relationships with the local micro-environmental patterns captured on H&E images.

### 2.6 Star-Motifs as potential prognostic markers

To dissect the prognostic associations of individual Star-Motif frequencies within tumor types, we correlated each motif’s frequency with OS and progression-free interval (PFI) (Methods). Using a single motif’s frequency to model risk, we computed c-index and checked its 95% confidence interval (CI). With this approach, c-index values above 0.5 indicate a link to poorer prognosis, whereas below 0.5 suggest favorable outcomes. Additionally, we performed a simple median-based patient stratification (high vs. low motif frequency) to see whether each motif alone could significantly separate patients into distinct risk groups (log-rank test, bootstrap-corrected *p* − *value <* 0.05). We present Star-Motifs with significant c-index CIs or significant stratifications in at least one cancer type in Fig. 6D and associated Kaplan–Meier plots for statistically significant patient stratifications in Figs.6E and S12-13. Together, our analyses revealed that several Star-Motifs correlate with good or poor prognosis, and simply splitting patients into high and low frequency groups result in significant risk stratification.

Next, we asked whether Star-Motif frequencies can be used to predict clinical outcomes. We used the patient-level vector representations, describing each patient as a composition of the Star-Motifs, as input to a classical machine learning model (Random Survival Forests - RSF) to predict overall survival (OS). We performed separate experiments for different input combinations: NEO Star-Motif frequencies, INF Star-Motif frequencies, and both combined (NEO+INF Star-Motif frequencies). We used a 5-fold cross validation over the data of each cancer type and reported mean (+-std) concordance index (c-index) values (Fig.6A). The performances were on par or better with those obtained by Chen *et al.* [16] using DL (Fig.6A). Moreover, RSF models demonstrated statistical significance in binary patient stratification with all input combinations in LUAD, with NEO+INF Star-Motif frequencies as input in BRCA, and with INF Star-Motif frequencies in CRC (Figs.6B and S9). We further plotted the SHAP (SHapley Additive exPlanations) values which can be used to interpret which motifs contributed most to model predictions (Figs. 6C,S10,S11) and observed that high or low feature values (high or low frequencies) of various motifs were associated with good or poor survival predictions.

Several motifs enriched in inflammatory cells (NEO-6,9,10,11,13; INF-2,8,9,10,11) demonstrated significant links to survival. For OS, some correlated with better outcomes (c-index CI *<* 0.5) in BRCA (NEO-13, INF-2) and LUAD (NEO-9). Moreover, several of these motifs stratified patients into distinct high- and low-risk groups, notably NEO-13 and INF-8 in BRCA, and NEO-11 and INF-8 in LUAD. Looking at PFI, NEO-10,11,13 and INF-2,8 aligned with better prognosis (c-index CI *<* 0.5) in BRCA, while NEO-9,11 were favorable in LUAD, and NEO-6 in LUSC. Risk stratification was similarly achieved in BRCA (NEO-13, INF-2,10), CRC (NEO-9), LUAD (NEO-11), and LUSC (NEO-6). By contrast, many of these same inflammatory motifs (NEO-10,11,13; INF-2,8,9,11) were detrimental in PRAD (c-index CI *>* 0.5), indicating that the prognostic impact of immune-enriched microenvironments may be highly context-dependent. Besides inflammatory-enriched motifs, we noted additional consistent trends. NEO-1, NEO-2 and INF-12, showing closer representation of neoplastic and connective cells compared to other cell types, were linked to poorer PFI (c-index CI *>* 0.5) in CRC and yielded significant log-rank tests. Conversely, motifs with characteristic enrichment of non-neoplastic epithelium cells (NEO-5, NEO-14, INF-16) correlated with improved PFI (c-index CI *<* 0.5) in BRCA, with NEO-14 and INF-16 also splitting patients into high- and low-risk categories. Meanwhile, NEO-0 and INF-3, each representing predominantly tumor and necrotic cell enrichment with low abundance of other cell types—were associated with worse PFI in BRCA and significant patient stratifications. Overall, these findings suggest that Star-Motifs capture various spatial patterns associated with patient survival and disease progression.

**Fig. 6.**
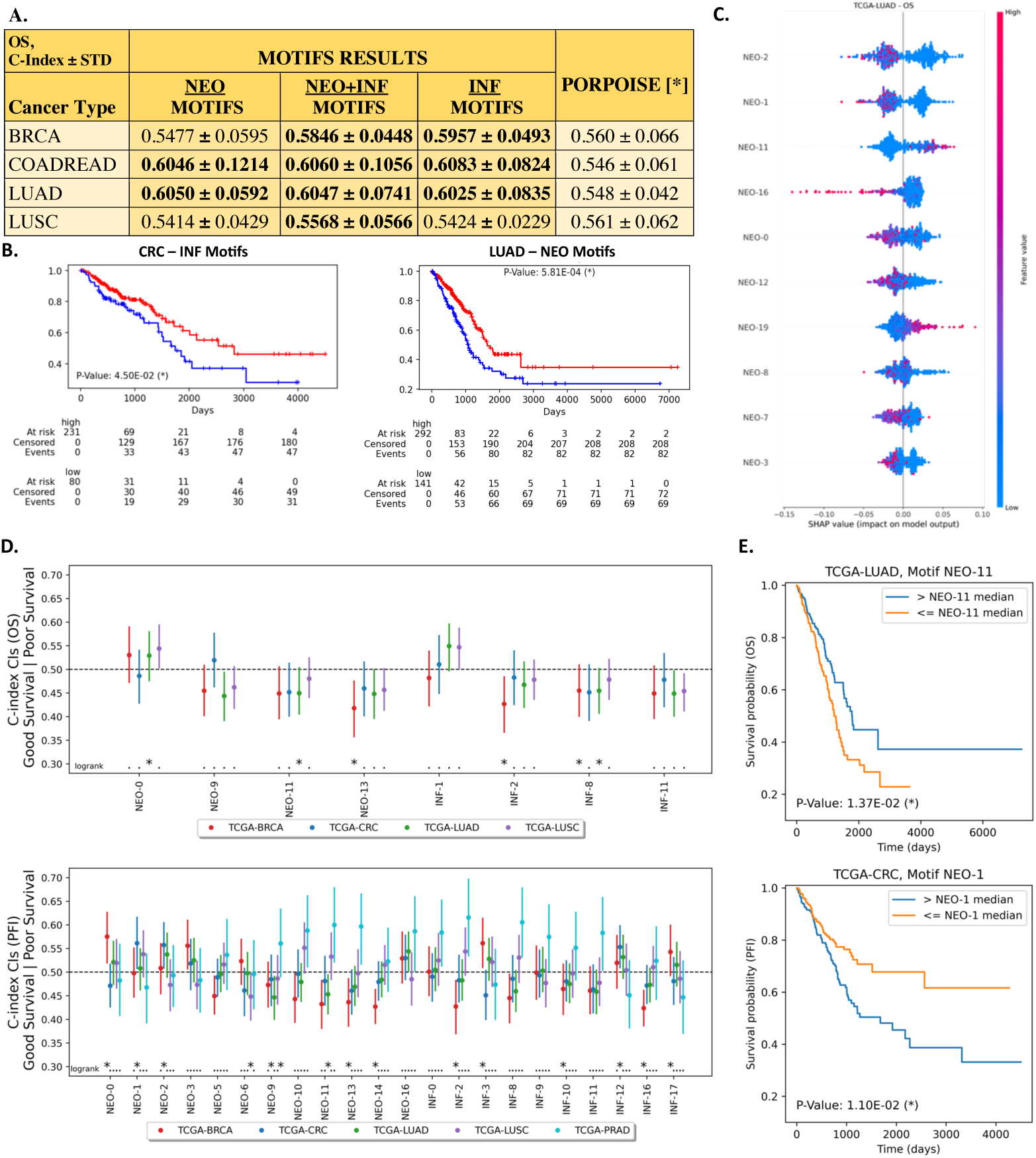
Star-motifs are predictive of survival. **A.** Results table of the Overall Survival (OS) prediction experiments with different combinations of Star-Motif frequencies as input to RSF models, mean(+-std) C-index values over 5-fold cross-validation are reported along with the results of the DL model of Chen *et al.* [16]. **B.** Kaplan-Meier analysis of stratification of low (blue) and high (red) survival patients via RSF model outputs on test sets. Plots show the experiments on CRC with INF motif frequencies as input and on LUAD with NEO motif frequencies as input. Log-rank test was used to test for statistical significance in survival distributions between patient groups (\**p <* 0.05). **C.** SHAP (SHapley Additive exPlanations) plot visualizing the contributions of Star-Motifs in the model predictions, showing the top ten contributing the most (for OS prediction on LUAD, with NEO motifs as input). A positive SHAP value (on the right side) is associated with better survival, while colors indicate the feature value, being high (red) or low (blue), associated with the SHAP scores. **D. Survival analysis of individual motifs.** Plots show C-index bootstrap confidence intervals for survival analysis of each motif and cancer type, for overall survival (top) and progression free interval (bottom); values below 0.5 indicate that a higher frequency of the motif favors longer survival (good outcome). Log-rank test results (if a motif can be used to stratify the high and low-risk groups) are also shown at the bottom with * if statistically significant (bootstrap corrected p value, 2 ∗ *p_median_*). **E.** Exemplary plots showing Kap1l6an–Meier curves for OS and PFI according to patient stratifications using the median values of individual motif frequencies (NEO-11, NEO-1) as the threshold, together with corresponding bootstrap corrected log-rank p-values.

### 2.7 Star-Motifs can predict clinically relevant targets

**Fig. 7.**
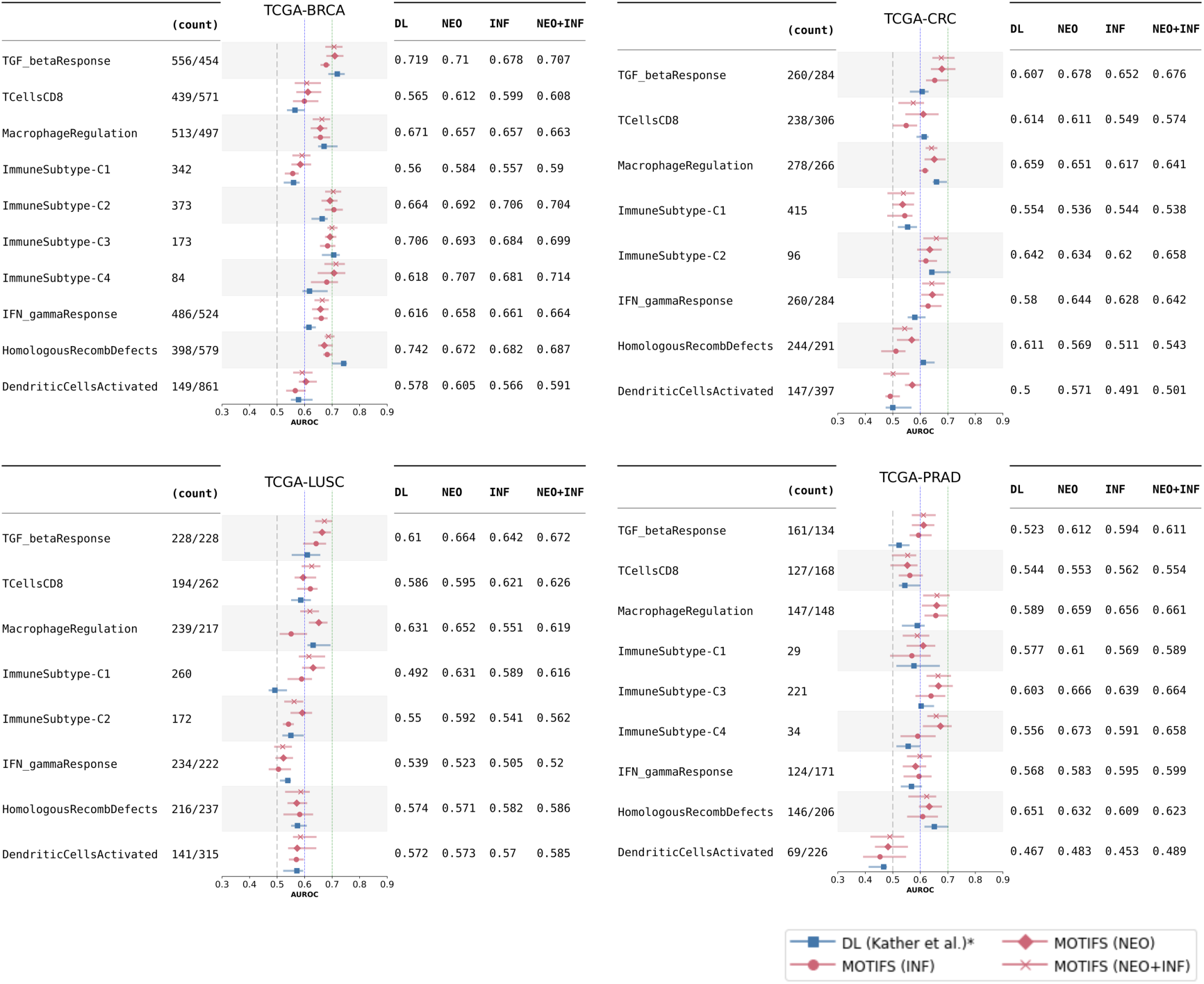
Mean AUROC values for predicting selected immune-related clinically relevant targets across four cancer cohorts (BRCA, CRC, LUSC, PRAD), demonstrating that random forest models using patient-level Star-Motif frequency vectors (NEO, INF, or both combined) as input can match or exceed DL performance [17]. Each subplot corresponds to one cancer type.

We evaluated the capability of *Star-Motifs* to predict clinically relevant alterations. For this, we performed extensive experiments to predict a wide range of targets such as molecular tumor subtypes, gene expression signatures and standard pathology biomarkers (in our standard/signature/subtype prediction experiments) and genetic mutations (in our mutation prediction experiments). These targets are used by Kather *et al.*[17] to investigate if these alterations leave a sufficient footprint in histomorphology to be inferred from histology images alone using a DL model. Similarly, we investigated if they can be inferred by only using information on different spatial organization patterns in each patient’s data, as summarized by their Star-Motif frequencies. We followed a similar experimental pipeline and adopted a 3-fold cross-validation strategy. We trained a classical machine learning model (i.e. random forests) for each experiment (see Methods) and reported the area under the receiver operating curve (AUROC) values on the validation sets. As input, we used the Star-Motif frequencies calculated on the patient level and performed separate experiments for different input combinations: NEO Star-Motif frequencies, INF Star-Motif frequencies, and both of them together (NEO+INF Star-Motif frequencies).

In standard/signature/subtype prediction experiments, as expected, Star-Motifs were not predictive for every phenotype in each cancer type (hold-out AUROC*<*0.6; see Figs. 11-18 for all results in each cancer type). However, they were predictive of various targets, especially on BRCA, CRC, LUSC, and PRAD cohorts. Surprisingly, on some tasks, the performances were comparable or better (±0.01 AUROC or better, macro-average AUROC for multi-class targets) to that achieved by black-box DL models [17]. With NEO motifs as input: 15 out of 16 different targets were predictable on BRCA, 6 out of 15 on CRC, 3 out of 10 on LUSC, and 8 out of 10 on PRAD. Some of them (6, 5, 3, and 4 of them, in the same order) had performances on par or better compared to the reference DL method [17]. The performances that were comparable or better were mostly observed on targets related to immune-related signatures (Fig. 7). Interestingly, it is observed by Kather *et al.*[17] that these immune-related targets had much higher predictability by DL compared to other targets they have investigated. To give more detail, binary prediction of high or low TGF-*β* response, using NEO motifs as input, showed comparable or better performances (AUROC values of: 0.710, 0.678, 0.664, 0.612; on BRCA, CRC, LUSC, PRAD); compared to the reported DL performances (AUROC values of:0.719, 0.607, 0.610, 0.523; in the same order). Similarly, binary prediction of high or low Macrophage Regulation showed comparable or better performances (AUROC values of: 0.663, 0.651, 0.652, 0.659; on BRCA, CRC, LUSC, PRAD); compared to the reported DL performances (AUROC values of: 0.671, 0.659, 0.631, 0.589; in the same order). Similar results are observed for the prediction of TCellsCD8 (BRCA, CRC, LUSC), Immune Subtypes (BRCA, CRC, PRAD), IFN gammaResponse (BRCA, CRC), DendriticCellsActivated (BRCA), Stemness-RNABased (CRC), Pan GynClusters (BRCA), WoundHealing (PRAD), Proliferation (LUSC). We should note that motifs did not perform well on lung adenocarcinoma dataset (LUAD), with only the MacrophageRegulation task having a predictive performance (AUROC≥0.6).

In mutation prediction experiments (Fig. S19), while motifs were not predictive for any mutation on LUSC and LUAD (AUROC*<*0.6); 3 out of 8 mutations were predictable on BRCA, 8 out of 18 on CRC, and 1 out of 3 on PRAD. From these, prediction performances for MAP3K1 and PIK3CA on BRCA dataset; FAT1, BRAF, APC, AMER1 on CRC dataset; and TTN on PRAD dataset were on par or better compared to DL. We observed some interesting results. For example, in AMER1 mutation prediction on CRC, the performances of the proposed method were relatively very high, especially with (NEO+INF) motifs as input (AUROC of 0.641) compared to DL (AUROC of 0.512). Similarly, in TTN mutation prediction on PRAD, motifs showed impressive performance (AUROC of 0.682 with NEO+INF), compared to the AUROC of 0.567 of the comparison DL method [17], and even comparable to the reported result (AUROC of 0.658) on the same dataset of a recent foundational DL model [44] which was specially trained on a very large dataset.

Overall, these findings showed the potential of *Star-Motifs* for inferring clinically relevant targets from routine H&E images. Additionally, this approach offers an interpretable link to these clinically relevant phenotypes by examining SHAP values from trained random forest models and identifying the motifs that contribute most strongly to predictions. For instance, in TGF-*β* classification on BRCA (Fig. S20), several connective cell enriched motifs (NEO-18, NEO-1, NEO-8, NEO-2, NEO-10) were linked to high TGF-*β* status, consistent with the known role of fibroblasts in TGF-*β* signaling. Also, although performance on certain phenotypes did not match that of deep learning models, our approach demonstrated noteworthy predictive power, such as an AUROC of 0.718 for estrogen receptor status and 0.651 for progesterone receptor status in breast cancer, showing promise for motif-based analyses.

## 3 Discussion

In this work, we introduced the Star-Motifs Framework to identify and quantify higher-order spatial organization patterns surrounding single cells and applied it to 3,105 diagnostic H&E whole-slide images across five cancer types (BRCA, CRC, LUAD, LUSC, PRAD). We first segmented 1.8 billion cells using Hover-Net. Then, by using our proposed *Star-Environment* representation, we encoded the spatial arrangement of multiple cell types surrounding individual neoplastic or inflammatory cells. By clustering Star-Environments, we extracted interpretable *Star-Motifs*, yielding 20 “NEO Star-Motifs” (Star-Motifs around neoplastic cells) and 18 “INF Star-Motifs” (Star-Motifs around inflammatory cells). These motifs spanned a broad spectrum of higher-order relationships in the local microenvironments. Our results demonstrated that frequencies of these motifs in patient’s data reflect a variety of TME characteristics, including tumor-infiltrating lymphocytes (TIL) densities, immune expression signatures, immune subtypes, and molecular estimates of different immune cells. We found that many motifs were individually associated with survival (OS, PFI) and can be used to stratify patients into distinct risk groups in multiple cancer types. We further demonstrated the predictive utility of Star-Motif frequencies for survival and various clinically relevant alterations, showing performance on par with or surpassing certain deep learning (DL) approaches on multiple tasks. Collectively, our observations demonstrated that Star-Motifs extracted directly from H&E-stained slides capture biologically meaningful and prognostically relevant local microenvironment patterns, provide insights into the functional heterogeneity within the TME, and can be used for interpretable prediction of clinical outcomes and some clinically relevant alterations.

Our preliminary findings presented here suggest that Star-Motifs could play a functional role in the TME. Our approach offers a set of interpretable features, the Star-Motif frequencies, that summarize local spatial organization patterns in histology images of patients. By linking these to established image, methylation, and gene expression based measures (e.g. TIL regional fraction, leukocyte fraction, proliferation, immune expression signatures, immune subtypes), we observed that cells classified under the same morphological label—neoplastic or inflammatory) can display different relationships with these different TME characteristics depending on their local microenvironment. For instance, motifs enriched in inflammatory cells showed positive correlations with TIL fractions and signatures related to immune response (including Macrophage Regulation, IFN-*γ*, and Lymphocyte Infiltration), while motifs enriched in fibroblasts were linked to TGF-*β* signaling, in agreement with the literature suggesting its role to drive stromal activation and immunosuppression[39, 40]. Furthermore, our integration of molecular immune cell estimates revealed that different immune cell subsets (e.g. T-cell or macrophage subsets) may have distinct associations with particular Star-Motifs, suggesting that variations in spatial patterns among histologically similar cells may reflect differences in functional states. While some of these functional state changes known to be associated with changes in the the local microenvironment, such as TAMs [43], earlier work on histology images has not reflected such heterogeneity. For example, Zhao et al.[34] clustered inflammatory cells on breast cancer tissue images using pair-wise graph based features, and reported that abundance of four of their clusters showed significant positive association with abundance of a variety of immune cell subsets (including CD8+, CD4+ memory activated, mast cells, and macrophages M0, M1, and M2). In contrast, our Star-Motif approach revealed more diverse associations, for example some motifs showed opposite associations with the pro-inflammatory M1 subtype and the tumor-promoting M2 subtype.

In many solid tumors, high TIL density has been associated with favorable patient outcomes [26, 27, 45]. In line with these, we observed that certain inflammatory-enriched motifs were associated with favorable overall survival and progression-free interval across multiple tumor types. For instance, inflammatory-enriched NEO-13, which demonstrated the strongest correlations with lymphocytes and CD8+ T cells in BRCA among NEO motifs, is associated with better OS and PFI and can stratify patients into risk groups for both OS and PFI. Yet, these same inflammatory-enriched patterns associated with poorer PFI in PRAD, which might be reflecting context-dependent roles of immune infiltration. Beyond inflammatory motifs, some neoplastic- and fibroblast-rich motifs were correlated with poorer PFI in CRC, whereas some motifs with non-neoplastic epithelial cell enrichment corresponded to better survival outcomes in BRCA. Collectively, these findings suggested that the Star-Motifs might capture a range of spatial patterns associated with patient survival and disease progression.

Finally, we demonstrated the predictive utility of Star-Motif frequencies as input to classical machine learning models (random survival forests for prognosis and random forests for other tasks), showing performance on par with or surpassing deep learning methods [16, 17], both in survival prediction and in classifying immune-related targets. We evaluated the predictive performance of three Star-motif sets (i.e., NEO, INF, and a combined NEO+INF set) as input features to our predictive models. Performances varied across tasks and cancer types; in certain scenarios, the NEO motifs outperformed the INF motifs and vice versa. Notably, the combined NEO+INF set did not consistently yield superior results, likely due to the increased dimensionality and redundant information, which may dilute the impact of the most informative features and hinder performance.

DL approaches have shown us that H&E stained histology images contain relevant information to identify patterns (possibly unknown to us but detectable by DL) related to many characteristics of the tissue (such as: patient’s survival [16, 41]; genetic mutations, molecular tumor subtypes, gene expression signatures, biomarkers [17–19]; drug sensitivities [20]; transcriptomic states [21]; and, intratumoral heterogeneity [46]) by training DL models in a supervised manner or by unsupervised clustering of features extracted from DL models. However, these DL approaches are considered to be “black boxes” [23–25], offering very limited interpretability with post-hoc analysis of high-attention regions of the model predictions or representative regions of clusters. Nevertheless, these DL studies emphasized the potential of gaining new insights, forming new hypotheses, and discovering new human-interpretable biomarkers by tracing these tasks (e.g., molecular alterations) back to specific regions and interrogating the cellular compositions in these regions. For this, they analyzed the cellular compositions in different clusters [41, 46] or high-attention regions of the DL model [16, 17, 20, 21], by using cell segmentation and classification models such as the Hover-Net [36], and they cross-linked the cellular composition of these regions to negative/positive label or low/high survival predictions. For example, with their post-hoc analyses, Chen *et al.* [16] observed that high attention regions in high survival patients had greater immune cell presence across all cancer types. Compared to these DL approaches, our proposed framework offers the advantage of direct interpretability by data-driven discovery of spatial organization patterns of cells and directly linking them to biological and clinical factors, enabling a clearer understanding of how complex spatial organization patterns contribute to tumor behavior.

Besides DL studies, traditional approaches performed spatial analyses of specific components of TME, especially the tumor-infiltrating lymphocytes (TILs) [26–30, 32]. These approaches used various, relatively simple, metrics such as: counts, densities, and statistical pair-wise correlation metrics of specific cell types; in tumor or other specifically segmented regions [12]. These efforts resulted in well-studied clinical biomarkers, such as the colon cancer ‘Immunoscore’ [26] metric quantifying the count and density of TILs, and ‘sTIL’ [29–31] quantifying the count and density of TILs in the stromal compartments. These studies provided useful tools and shed light on the effects of the spatial distribution of specifically chosen cell types in pre-selected areas with pre-prepared metrics. However, these methods do not make full use of the complexity of the spatial data that can be obtained from H&E images. Many different cell types along with their spatial locations can be extracted from H&E images. As we have shown here, a higher-order spatial analysis method that allows interrogation of the organization of different cell types helps us identify the patterns of cellular compositions beyond TILs and map them to clinical and genomic characteristics.

Diao *et al.* [42] extracted human-interpretable features (HIFs) from H&E images, demonstrating strong predictive power and interpretability. However, their approach necessitates extensive pathologist-provided annotations to train multiple tissue- and cell-level segmentation models, with each feature carefully engineered (e.g., densities, morphological measures). By contrast, our *Star-Motifs Framework* adopts an entirely data-driven pattern discovery process that relies on a single nuclei segmentation and classification pipeline, substantially reducing annotation overhead. In addition, while many of Diao *et al.*’s features focus on densities or pairwise relationships, our Star-Environment representations explicitly capture higher-order multi-cell-type co-localizations around each neoplastic or inflammatory cell. Clustering these Star-Environments yields “Star-motifs” that can be located and visualized directly on tissue sections, facilitating straightforward insights into potentially complex TME configurations. Consequently, *Star-Motifs* not only provides a powerful alternative to feature engineering pipelines that require large-scale annotation, but it also offers a more flexible and scalable approach to interrogate diverse cancer cohorts for biologically relevant higher-order spatial patterns.

Graph (network) based approaches, such as the method proposed by Zhao *et al.* [34], model spatial architecture by constructing adjacency cell graphs for each pair-wise cell-type combination and use topological features extracted from these graphs to cluster individual cells. While these methods can be powerful, they can be sensitive to how adjacency is defined (e.g., via short distance thresholds) and they require constructing a separate adjacency graph for each cell-type pair. By contrast, our *Star-Motifs Framework* encodes the multi-cell-type organization around each cell within a continuous polar-coordinate vector (“Star-Environment”) capturing both distance and angular relationships in a large radius, offering an alternative perspective and a more straightforward interpretability compared to topological features (e.g., clustering coefficient, harmonic centrality).

A major contribution of our work lies in the development of the *Star-Motifs* framework itself. We provide a conceptual framework which provides a new lens for interrogating the spatial behavior of different cancer’s TME in histology images. The *Star-Environment* representation, as one of the key components of the framework we proposed here, is obtained by constructing multiple star-convex polygons, each representing the spatial arrangement of a different cell type surrounding the single cell at the center. These star-convex polygons can be imagined as the boundaries of the closest cells from different cell types surrounding the center cell. This type of data-driven representation (i.e., a star-convex polygon) have been used previously in the literature for describing object shapes and clustering them in natural images [47] and in fluorescence images [48]. These approaches, very differently than us, represented object shapes using a single polygon representation. To our knowledge, there have not been any strategy to use any similar type of representation to describe spatial organization patterns.

Our approach enables us to represent and systematically dissect complex, multicell-type spatial organizations in a straightforward, data-driven manner, suitable for large-scale datasets. Further, our unsupervised discovery of Star-Motifs ensures that we are not constraining the analysis to predefined regions, cell subtypes, or pair-wise metrics. Because motifs represent different spatial arrangements around cells, the motif frequencies offer an interpretable, low-dimensional patient descriptor, providing a holistic picture of TME architecture. Additionally, visualizing the motif maps on whole-slide images provide spatially localized insights into tumor architecture, which can guide both hypothesis generation in research and risk assessment in clinical settings.

The higher-order data-driven patterns extracted by the framework allowed us to detect context-dependent associations of these spatial organization patterns across different tumor types; for instance, we observed distinct TGF-*β* correlations for the same motif in colorectal versus breast cancers. Such insights may inform a deeper understanding of how spatial arrangement of multiple cell types collectively shape disease progression. Notably, the motifs’ interpretable nature enhances transparency, and might aid clinician confidence and the adoption in clinical workflows. Associations of Star-Motifs to prognosis showed the potential of Star-Motif frequencies as practical features for risk stratification and biomarker discovery from routine H&E slides. Also, as studies [7, 49] have shown that the immune contexture and its spatial relationship with the tumor cells has predictive value for immunotherapy response, our framework could support decisions on immunotherapies, targeted therapies, or combination treatments.

Despite its strengths, our approach has certain limitations. Star-Motif extraction relies on cell segmentation and classification; and errors at this stage may propagate into downstream analyses. We used Hover-Net [36], a model widely adopted in computational pathology; for instance, Zhao *et al.* [34] reported near-perfect correlation between Hover-Net cell classifications and CODEX on two samples (Spearman coefficients: 0.99 for tumor cells, 0.96 for inflammatory cells, and 0.95 for stroma cells). We also excluded 55 WSIs exhibiting substantial artifacts (e.g., folds or blurry regions) to minimize segmentation errors. Given that our framework integrates data from a large number of cells per patient, we anticipate that occasional misclassifications may have only a modest impact on patient-level Star-Motif frequency vectors. Additionally, though our pan-cancer dataset from TCGA is large, further validation on independent, multi-institutional cohorts would be valuable to establish clinical generalizability. Future directions might also extend Star-Motifs to additional cancer types.

From a methodological standpoint, the current Star-Environment representation, which relies on star-convex polygon shape descriptors, is directional and not rotation-invariant by construction. We address this by merging certain clusters based on their sensitivity to rotation, yielding rotation-robust TME features for analysis. Nevertheless, developing inherently rotation-invariant representations remains a promising extension; future work could explore alternative shape descriptors or learned rotation-invariant embeddings. While a large profiling radius, fine angular binning, and an empirically selected number of clusters were used in this study, systematic evaluation of these parameter choices would further clarify the sensitivity of the results.

Beyond the spatial organization captured here, integrating nuclear morphological metrics or complementary spatial omics data could yield richer biological insights. Finally, modeling the spatial relationships between motifs themselves, rather than only their frequencies, may reveal organizational principles within the TME.

## 4 Methods

### 4.1 Ethics statement

This research study was conducted retrospectively using human subject data made available in open access by The Cancer Genome Atlas (TCGA) Research Network (https://www.cancer.gov/tcga). As such, ethical approval was not required as confirmed by the license attached with the open-access data.

### 4.2 Data

We collected 3,160 H&E-stained diagnostic whole-slide images (WSIs) across six TCGA cancer types: breast invasive carcinoma (BRCA, 1106 WSIs), colon adenocarcinoma (COAD, 439), rectum adenocarcinoma (READ, 157 WSIs), lung adenocarcinoma (LUAD, 526 WSIs), lung squamous cell carcinoma (LUSC, 510 WSIs), and prostate adenocarcinoma (PRAD, 422 WSIs). Following previous studies [16, 17], we grouped the data from the gastrointestinal tract (COAD and READ cancer types) together, forming the combined colorectal cancer (CRC) cohort used throughout the study. After the nuclear segmentation and classification step, if a WSI showed large unsegmented areas or inconsistent nuclear classification, it was manually inspected and excluded. After excluding 55 WSIs with substantial artifacts that impeded reliable segmentation (e.g., tissue folding, out of focus/blurry scanning), the final dataset comprised 3,105 WSIs from 2,892 patients.

In correlation analyses, we used data compiled and published by Thorsson *et al.* [5]. Specifically, we used:

- TIL regional fraction: the spatial fraction of tumor regions with TILs, estimated from TCGA H&E images using deep learning-based lymphocyte classification (Saltz *et al.*[27])
- The leukocyte fraction (LF): representing leukocyte content estimated using Illumina Infinium DNA methylation platform arrays
- Stromal fraction (SF): representing the total non-tumor cellular component
- Proliferation
- Immune cellular fraction estimates: the relative fraction of immune cell types estimated on TCGA RNASeq data. In our analyses we included:

– Mast cells (Mast cells resting + Mast cells activated)
– Macrophages M0
– Macrophages M1
– Macrophages M2
– Lymphocytes (B cells naive + B cells memory + T cells CD4 naive + T cells CD4 memory resting + T cells CD4 memory activated + T cells follicular helper + T cells regulatory (Tregs) + T cells gamma delta + T cells CD8 + NK cells resting + NK cells activated + Plasma cells)
– T cells CD8
– T cells CD4 memory activated
- Characteristic immune expression signature scores: wound healing, TGF-*β* response, IFN-*γ* response, macrophage regulation, lymphocyte infiltration
- Immune subtypes: six groups (C1-C6) characterizing intratumoral immune states, obtained by clustering the immune expression signature scores

Survival data was collected from the TCGA Pan-Cancer Clinical Data Resource (TCGA-CDR) [50] and the values of OS, OS.time, PFI, and PFI.time were used. Following their recommendations [50], we have only used the PFI endpoint for the TCGA-PRAD cancer type, while other cancer-types were included for both endpoints.

For predictive analyses, we used the clinically relevant targets compiled and published by Kather *et al.* [17].

### 4.3 Nuclear segmentation and classification

In all WSIs, tissue regions were segmented from the background using Otsu thresholding [51]. Then, we employed the Hover-Net model [36] (trained on the PanNuke dataset [52]) on these tissue regions, to segment individual nuclei and classify each nucleus into one of five categories:

- Neoplastic (tumor) epithelium
- Connective (including fibroblasts and endothelial cells)
- Inflammatory (including lymphocytes, leukocytes, and macrophages)
- Non-neoplastic (normal) epithelium
- Dead (necrotic) nuclei

All segmentation and classification computations were performed on each WSI at at a spatial resolution of 0.50 microns-per-pixel (MPP) using multiple *NVIDIA A100 GPUs*. A total of *1,833,975,602* nuclei were ultimately segmented across all cancer types in the final dataset.

### 4.4 Star-Environment profiling

Star-convex polygons have been applied to represent object shapes in other image analysis applications [47]. Here, we adapted this idea to capture the surrounding multi-cell-type local microenvironment for each cell, rather than the shape of a single object. We developed the “Star-Environment” representation, which is obtained by constructing multiple star-convex polygons, each representing the spatial arrangement of a different cell type surrounding a cell at the center.

Specifically, for a given cell *C* of interest, we consider all neighboring cells within a defined maximum radius *r*_max_. We then convert each neighbor’s Cartesian position into polar coordinates (*r, θ*) centered on *C* with the angular domain discretized into N bins. For each of five cell-type classes (neoplastic, inflammatory, connective, dead, and non-neoplastic), we record the radial distance *r* to the nearest cell of that type at every angular bin (*θ_i_*, i = 1,2,…,N). If no cell of a given type lies within *r*_max_ in a particular bin, that bin is assigned the maximum radius. We end up with five *r_celltype_* vectors (*r_celltype_* = [*r*_1_*, r*_2_*, r*_3_*, …, r*_N_]), one for each different cell type, each capturing the radial distribution pattern across angular directions. Concatenating these five *r_celltype_* vectors yields a (5 × N)-dimensional vector for each cell:

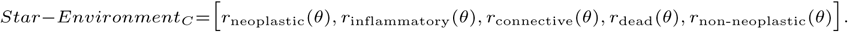

This Star-Environment representation thus captures how multiple cell types are radially arranged around a center cell. We computed these representations for all neoplastic and inflammatory cells in each WSI.

**Choice of angular binning and radius thresholds.** The maximum radius and angular binning were chosen to encompass all plausible cell–cell interactions and to maintain high directional resolution. Specifically, the maximum radius was set to twice the established effective cell-to-cell communication distance of 250*µm* [37] (*r*_max_ = 500*µm*), and angular binning was defined at one-degree resolution (N = 360 bins) to capture fine-grained patterns, with dimensionality reduction handled by singular value decomposition (SVD), as described in the following subsection. Both parameters were intentionally set to large values to avoid constraining the representation.

### 4.5 Data-driven extraction of Star-Motifs

The identification of the Star-Motifs, recurrent surrounding spatial organization patterns, is done through a series of steps (i.e., unsupervised clustering after dimensionality reduction) on a dataset of extracted Star-Environment vectors. To perform these steps, we decided to use the same number of Star-Environment vectors from each WSI to avoid any potential biases towards WSIs with higher cellularity. To this end, we sampled 2,000 neoplastic cell Star-Environments and 1,000 inflammatory cell Star-Environments from each WSI and assembled two large matrices **A_NEO_** (with shape 6,210,000 X 1,800) and **A_INF_** (with shape 3,105,000 X 1,800). Following steps were performed separately on *A_NEO_* and *A_INF_*, resulting in two sets of Star-Motifs: *NEO Star-Motifs* and *INF Star-Motifs*.

#### 4.5.1 Dimensionality reduction via SVD

As each Star-Environment vector has high dimensionality (1800), we first performed dimensionality reduction via low-rank approximation. By doing so, we get to represent a Star-Environment vector by only using a small number of values. Emphasizing that the distances between vectors in the original space are equal to those between the corresponding vectors in the low-rank space, this low-rank approximation approach aids in performing clustering more reliably and more efficiently by alleviating the curse of dimensionality [47, 53].

Specifically, we performed truncated singular value decomposition (SVD) to achieve best rank-M approximation. To pick the number of components (M) which will optimize the total explained variance, we investigated the ratio of variance explained by each component, and detected the point of maximum curvature (a knee point) using the “kneedle” algorithm [54] (using the Python implementation of [55]). The M values are selected as 42 for both *A_NEO_* and *A_INF_* independently, with first M components capturing respectively 39.0% and 43.3% of the total amount of variation in the matrices.

#### 4.5.2 Clustering to identify Star-Motifs

We separately performed K-means clustering on the best rank-M approximations of *A_NEO_*and *A_INF_* . To choose the optimal number of clusters (k), we systematically evaluated cluster stability over a range of *k* ∈ [2, 32] using the “Cluster Validation by Prediction Strength” method of Tibshirani *et al.* [56], which is also used by Thorsson *et al.* [5]. This method provides a strategy for determining the largest number of reproducible clusters.

Specifically, for a given *k*, we randomly split the data into equal-sized training and test subsets, and trained independent k-means models on each subset. For each testset cluster, we then calculated the proportion of observation pairs in that cluster that are also assigned to the same cluster by the model trained on the training subset. The prediction strength is defined as the minimum of these proportions across all test-set clusters. We repeated this procedure five times with different train/test splits (reusing the same splits for each *k*). Following Tibshirani *et al.* [56], we selected the largest *k* whose mean prediction strength across these five runs exceeded 0.8, a threshold indicated by the authors as a good reproducibility criterion. As shown in Supplementary Fig. 21, the optimal cluster count was *k* = 20 for *A_INF_* and *k* = 22 for *A_NEO_*.

Next, we merged certain clusters based on rotation invariance. After initial inspection, we noted that some clusters appeared to be rotated variants of one another. To formalize this, we systematically rotated the Star-Environment vectors by multiples of 45° ([45, 90, 135, 180, 225, 270, 315]) and reassigned them to the closest centroid in each *k*-means model, examining confusion matrices to detect consistently “confused” clusters. In both NEO and INF analyses, three clusters were thus merged into a single Star-Motif.

In total, we arrived at 20 final NEO Star-Motifs (one comprised of three merged clusters) and 18 final INF Star-Motifs (again, one merged triplet).

### 4.6 Concentric enrichment analysis of Star-Motifs

To quantify how the cell distribution differs in the vicinity of each Star-Motif, we performed a “concentric enrichment” analysis (separately on the **A_NEO_** and **A_INF_** matrices, Fig. 2C, Supplementary Figs. 1–2). Each Star-Environment vector encodes the distance to the nearest cell of each type at 360 discrete angles (Section 4.4). For each distance threshold *d_i_* ∈ {25, 50, 125, 250, 500} *µ*m, we counted how many angles satisfy *r*_celltype_(*θ*) *< d_i_*: 1. Across all Star-Environments in the relevant matrix, and 2. Only those Star-Environments assigned to a particular Star-Motif in that matrix. If C*_di_* denotes the average count of a certain cell type appearing at distance *< d_i_* in all Star-Environments, and C^motif^ the corresponding average count in Star-Environments assigned to a particular Star-Motif, we define the log(fold-enrichment) as:

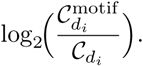

A positive log(fold-enrichment) thus indicates that the cell type is comparatively overrepresented at distance *d_i_* within that Star-Motif. By examining multiple thresholds, we gain a layered view of how various cell types cluster around each Star-Motif at different radii, highlighting distinct patterns of local microenvironmental organization.

### 4.7 Patient-level Star-Motif frequency vectors

To use in our analyses, we summarized each patient’s Star-Motif distributions by calculating the Star-Motif frequencies (proportions). To obtain these patient-level descriptors, for each neoplastic cell in each WSI, we have obtained the corresponding NEO Star-Motif label (i.e., which NEO Star-Motif each neoplastic cell’s environment exhibits) based on the nearest cluster centroid in the low-dimensional embedding, and similarly, for each inflammatory cell in each WSI, we have obtained the corresponding INF Star-Motif label. We then calculated, for each patient:

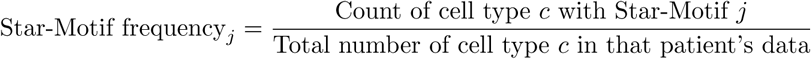

where *c* is {neoplastic, inflammatory} and *j* indexes the discovered Star-Motifs (e.g., 20 for NEO, 18 for INF). This yields a frequency vector of length 20 for NEO Star-Motifs or 18 for INF Star-Motifs per patient. In subsequent analyses, these Star-motif frequency vectors served as the primary input.

### 4.8 Correlations with tumor microenvironment measures

We correlated patient-level Star-Motif frequencies with established tumor microenvironment measures (e.g., TIL fractions, immune expression signatures, immune subtypes) compiled and published by Thorsson *et al.* [5]. Correlations were assessed using Kendall’s *τ*, and multiple comparisons were adjusted via the Benjamini–Hochberg procedure [57]. Corrected P-value ≤ 0.05 is considered significant.

### 4.9 Survival prediction

We assessed predictive power of Star-Motif frequency vectors for overall survival (OS). Patient’s Star-Motif frequency vectors were used as input to Random Survival Forest (RSF) model (using RandomSurvivalForest from sksurv library). For each cancer cohort, using the data splits of [16], we divided the data into 5-folds, we performed 5-fold cross-validation and reported the mean Harrel’s concordance index (c-index) on validation sets. At each cross-validation fold, input values are normalized according to the discovery sets by removing the mean and scaling to unit variance. Also, the optimal model parameters are selected on discovery sets at each cross-validation fold, using GridSearchCV (from Python Scikit-learn library). With this grid search approach, at each fold, the discovery set was further divided into 3 folds and parameter selection was performed over a parameter search space (n estimators:[2000], min samples split:[6], max depth:[10, 20, 30, None], max samples:[0.25, 0.5, 0.75, 1], min samples leaf: [3, 4, 5, 6, 7, 8, 10], max features: [2, 4, 5, 6, 7, 8, 9]).

Kaplan–Meier curves were generated by splitting patients into high-risk and low-risk groups based on model outputs. We selected the optimal cut-off values in the discovery sets and used these cut-off values in the test sets to divide the subjects in the test sets into two groups. We selected an optimal cut-off value as the value that yields the maximum log-rank test statistic out of a series of candidate values generated across the range of predicted risk scores, while excluding the lowest and highest 10% of values to ensure that both resulting groups were sufficiently large. To plot the Kaplan-Meier curves, we aggregated the risk groups from all test sets and plotted them against their survival time. We used log-rank tests to measure if the difference of two survival distributions is statistically significant (P-value *<* 0.05).

For model interpretability, we calculated SHAP (SHapley Additive exPlanations) values (using shap.Explainer function) on the test sets to visualize the contribution of each Star-Motif in the RandomSurvivalForest outputs. Each plot shows, in order, the top ten Star-Motifs contributing the most. A positive SHAP value (on the right side) is associated with better survival, while colors indicate the feature value, being high (red) or low (blue), associated with the SHAP scores.

### 4.10 Prognostic associations of individual Star-Motifs

We evaluated the relationship between individual Star-Motif frequencies and clinical outcomes, focusing on overall survival (OS) and progression-free interval (PFI). To quantify these associations, we directly used the individual Star-Motif frequencies to model risk and calculated Harrel’s concordance index (c-index). With this approach, for an individual Star-Motif, c-index values above 0.5 indicate a correlation with poor prognosis, whereas intervals below 0.5 indicate an association with favorable outcomes. Additionally, for each motif, we checked, using log-rank test, if a simple binary stratification (i.e., using the the median value of individual Star-Motif frequencies in each cancer type as threshold for splitting) can differentiate patients into distinct risk groups. We used bootstrapping (for 1000 bootstrap resamples each) to calculate C-index confidence intervals (C-index CIs) and patient stratification test results and reported the mean C-index, 95% C-index CIs and bootstrap-corrected log-rank p values (2\**P_median_*).

### 4.11 Predicting clinically relevant targets

We assessed predictive power of Star-Motif frequency vectors for various clinically relevant features (e.g., gene expression signatures, molecular subtypes, mutation status) we used the targets compiled and used by Kather *et al.* [17]. Patient’s Star-Motif frequency vectors were used as input to random forest classification models (using RandomForestClassifier from sklearn library). Similar to Kather *et al.* [17], we adopted a 3-fold cross-validation strategy. To evaluate performance, we reported the mean area under the receiver operating characteristic curve (AUROC) values over a 10x bootstrapped experiment with a confidence interval representing lower and upper range. For binary prediction tasks, the performance for the positive labeled class is reported.

At each cross-validation fold, input values are normalized according to the discovery sets by removing the mean and scaling to unit variance. Also, we selected the optimal model parameters on discovery sets at each cross-validation fold using GridSearchCV (from Python Scikit-learn library). With this grid search approach, at each fold, the discovery set was further divided into 3 folds and parameter selection was performed based on the AUROC score (roc auc score function from the Scikitlearn, with parameters: average=‘macro’, and multi class=‘ovo’ for multi class targets) over a parameter search space (criterion: [gini, entropy, log loss], n estimators:[2000], min samples split:[6], class weight:[balanced], max depth:[5, 10, 20], max samples:[0.1, 0.25, 0.5, 0.75], min samples leaf:[3, 4, 6, 8, 12], max features:[2, 3, 4, 5, 6]).

For model interpretability, we calculated SHAP (SHapley Additive exPlanations) values (using shap.TreeExplainer function) on the test sets to visualize the contribution of each Star-Motif in the RandomForestClassifier outputs. The example plot shows, in order, the top ten Star-Motifs contributing the most. A positive SHAP value (on the right side) is associated with the positive label class, while colors indicate the feature value, being high (red) or low (blue), associated with the SHAP scores.

### 4.12 Implementation details

All analyses were performed in Python (version 3.8). Packages used include NumPy, SciPy, scikit-learn, Pandas, Lifelines.

Whole slide processing, tissue region extraction, and the Hover-Net cell instance segmentation and classification was implemented using the TIAToolbox library [58]. The trained Hover-Net deep learning model [36] on the PanNuke dataset [52] were available on (https://github.com/TissueImageAnalytics/tiatoolbox).

We implemented *Star-Environment Profiling* code using Numba and Cuda packages, optimizing it to be fast using parallelized GPU computing.

We used the Dask package which helped with scalability of computations and dimensionality reduction on large matrices. Specifically, we stored the sampled Star-Environment matrices (*A_INF_* and *A_NEO_*) as dask.array’s, and we used the dask.TruncatedSVD method which performs truncated singular value decomposition.

The “kneedle” algorithm [54] to select the optimal number of components for dimensionality reduction is applied using the Python implementation of [55].

Plotting and visualization were obtained using Seaborn, and Matplotlib. We also used Shap package for model interpretability where applicable.

Hover-Net inference and Star-Environment profiling were carried out on multiple NVIDIA A100 GPUs.

### 4.13 Data availability

All slide images, along with associated metadata, were retrieved from the NCI Imaging Data Commons (https://portal.imaging.datacommons.cancer.gov). The processed nuclear segmentation and classification results, along with the corresponding Star-Environment vectors and final Star-Motif labels, will be made publicly available upon publication. Additional requests should be directed to the corresponding author.

### 4.14 Code availability

Scripts to perform nuclei segmentation and classification and the trained Hover-Net model are available on github (https://github.com/TissueImageAnalytics/tiatoolbox).

Source code and documentation of all Python scripts used in the study will be made available upon publication.

For any further information, please contact the corresponding author.

**Supplementary information.** The article has accompanying supplementary files.

## Acknowledgements

G.N.G. would like to acknowledge the PhD studentship support from GlaxoSmithKline and the Department of Computer Science at the University of Warwick.

