## Supplemental Materials for "Star-Motifs: Revealing Single-Cell Spatiotypes from Routine Histology Using Star-Convex Neighborhoods"

**Supplemental Materials:** *Star-Motifs*

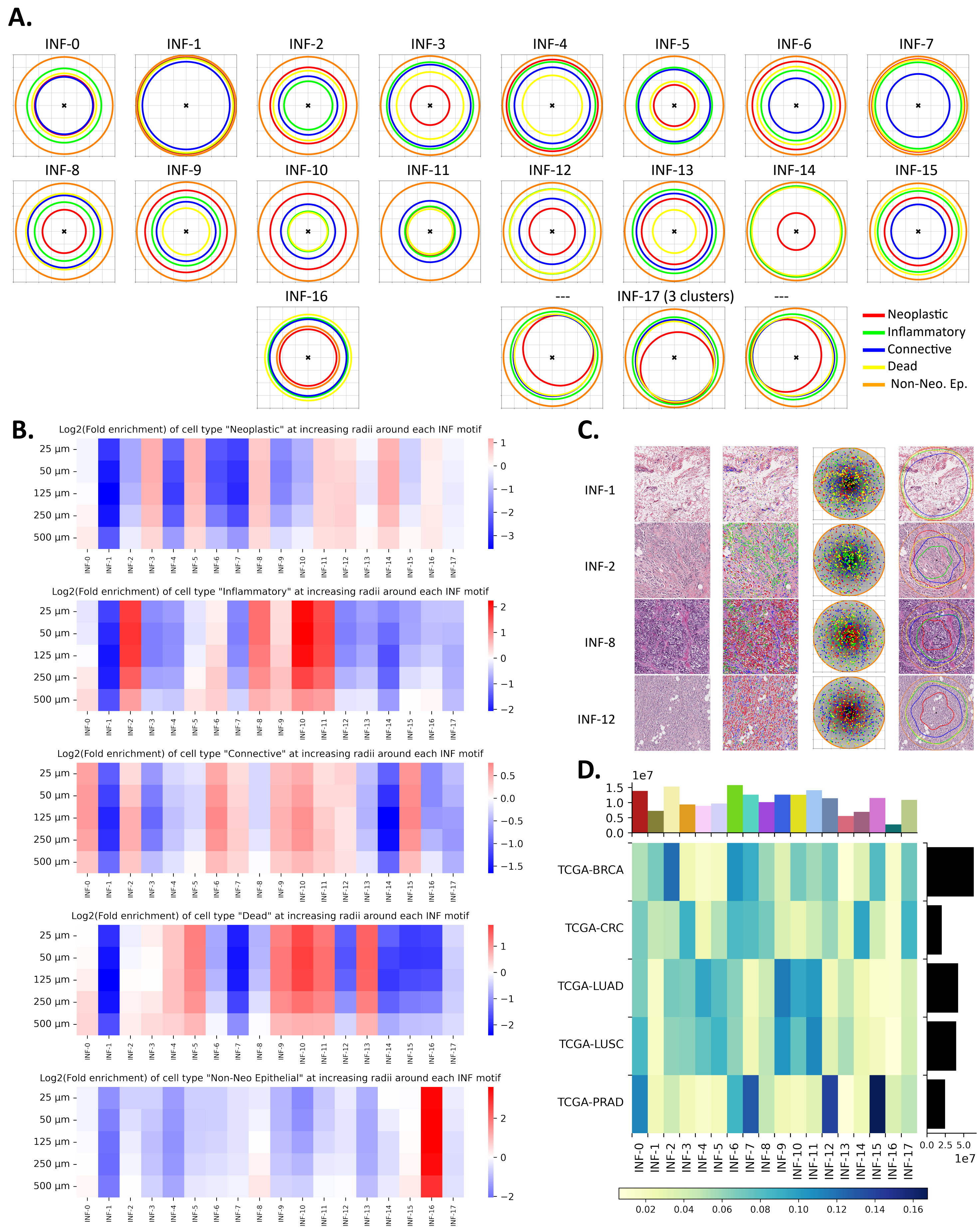


**S1: INF Star-Motifs A.** Extracted INF Star-Motifs are illustrated. Each plot in the figure shows the reconstructed cluster centre from the low-rank space to the original space. These Star-Motifs, which are obtained in a data-driven manner, represent the approximations of different higher-order patterns of different cell types surrounding a cell in the centre (with the centre cell being an inflammatory cell in INF Star-Motifs). **B.** The number of different types of cells surrounding each INF cell with increasing concentric window sizes. Showing log(fold enrichment) values of each cell type at each distance for each INF Star-Motif. **C.** Most representative Star-environments (closest to the cluster centres). Shows image patches with the cells having the most representative environment at the patch centre (column 1), corresponding nuclei segmentation (column 2), Star-environments before (column 3) and after (column 4, overlaid on image patches) low-rank transformation. **D.** INF Star-Motifs VS cancer type. Heatmap showing the fraction of cells in each of the eighteen INF Star-Motifs in each cancer type. Rows were normalised to sum to 1. Histograms show the total number of cells exhibiting each motif and in each cancer type. All inflammatory cells were included to compute the heatmap and the histograms.

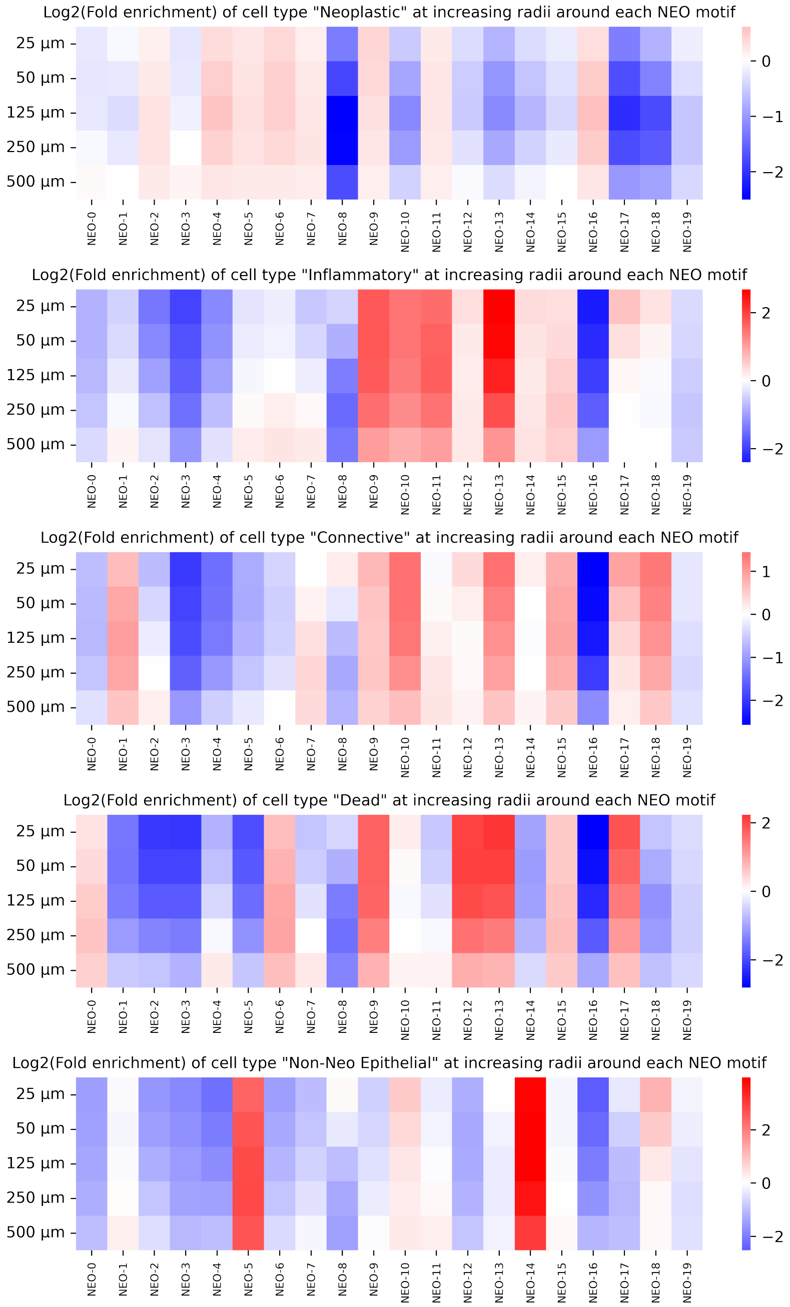


**S2:** The number of all different types of cells surrounding each neoplastic cell with increasing concentric window sizes. Showing log(fold enrichment) values of each cell type at each distance for each NEO motif.

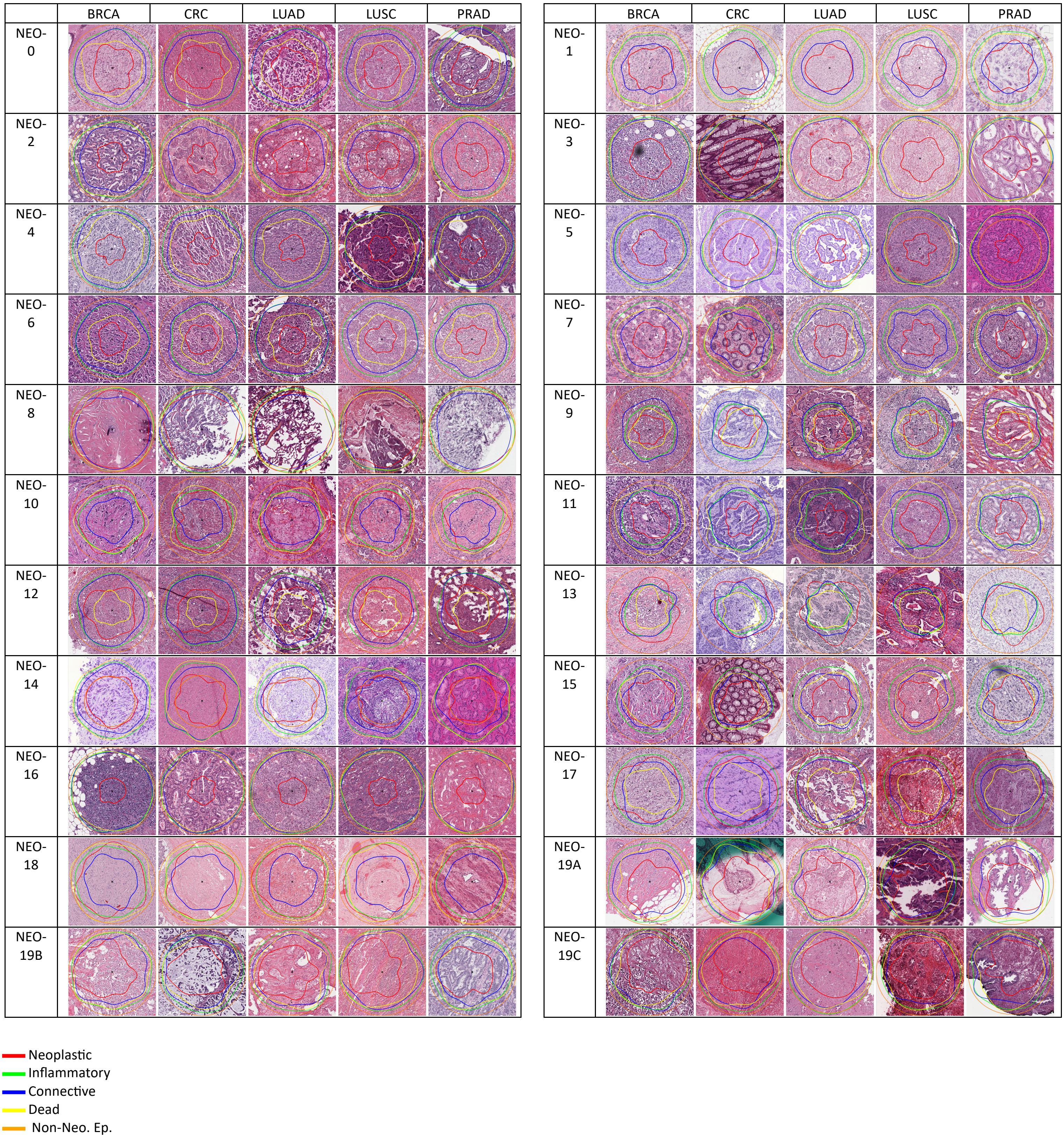


**S3:** Most representative Star-Environments of each NEO Star-Motif (closest to the cluster centres) from each cancer type. The table shows image patches with the cells having the most representative environment at the patch centre with their Star-environment approximations overlaid on the image patches.

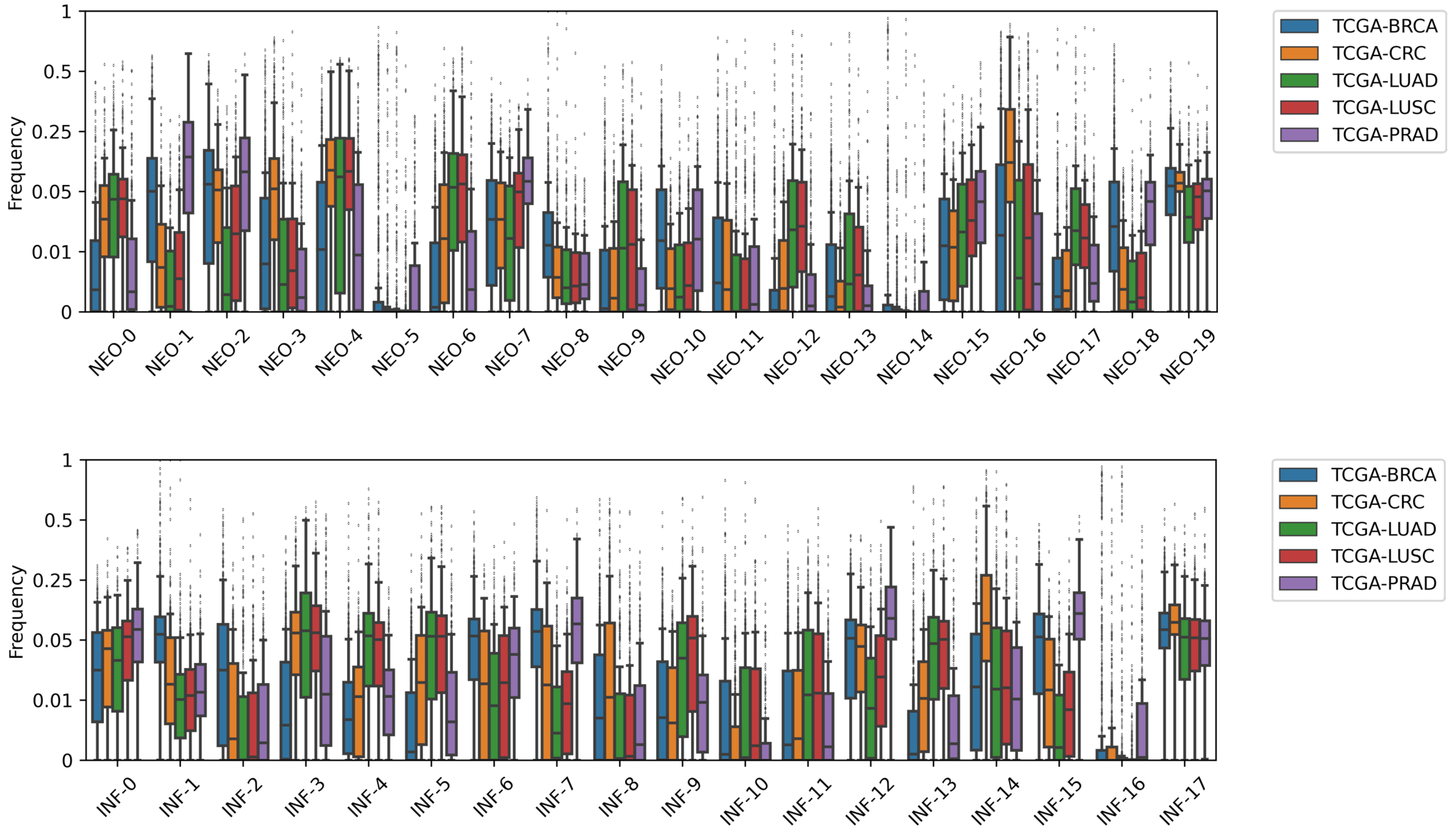


**S4:** Star-Motif frequency distributions of patients for NEO Star-Motifs (top) and INF Star-Motifs (bottom). Boxes indicate the quartile values of data distributions, and whiskers extend to data points within 1.5 times the interquartile range. Plots use a custom y-axis scale to enhance visibility.

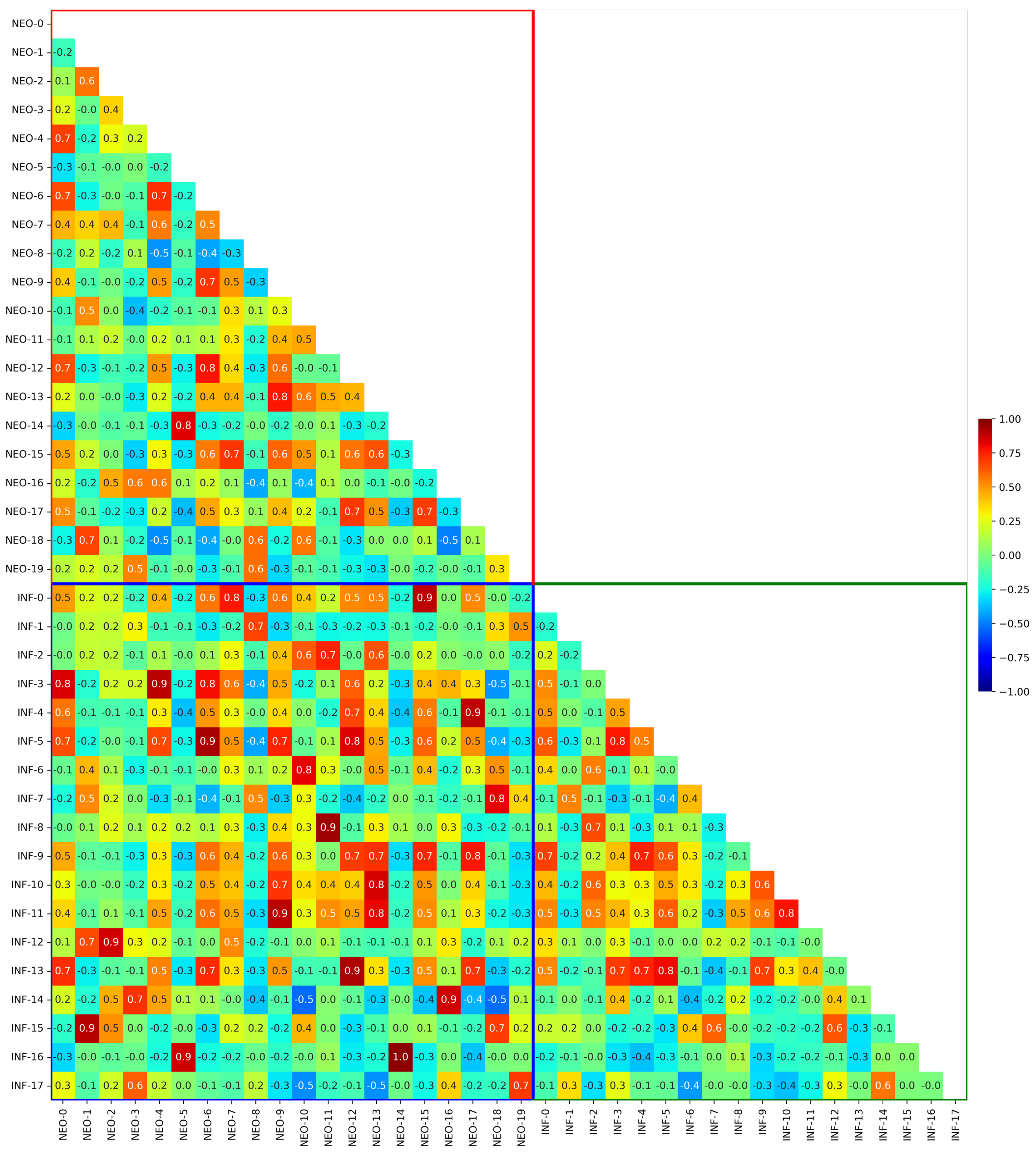

**S5:** Spearman correlations of different Star-Motif frequencies in patients' data. The figure contains cross-set correlations (INF vs NEO) inside the blue square and intra-set correlations (NEO vs NEO and INF vs INF) inside the red and green squares.

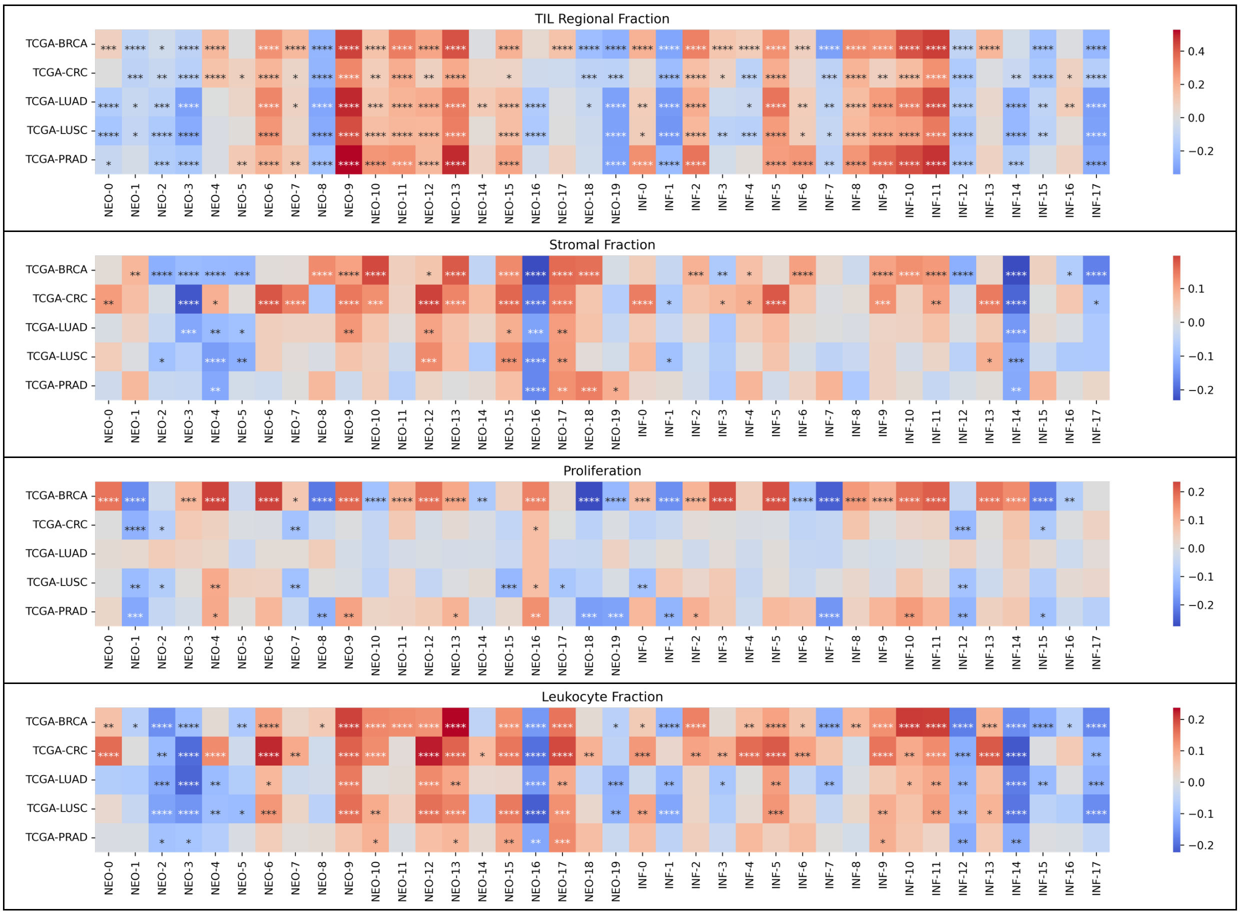


**S6:** Associations of Star-Motif frequencies with TME characteristics, across cohorts. Red and blue colours indicate the degree of association (Kendall's tau correlation coefficients, FDR BH correction). *P≤0.05, **P≤0.01, ***P≤0.001, ****P≤0.0001.

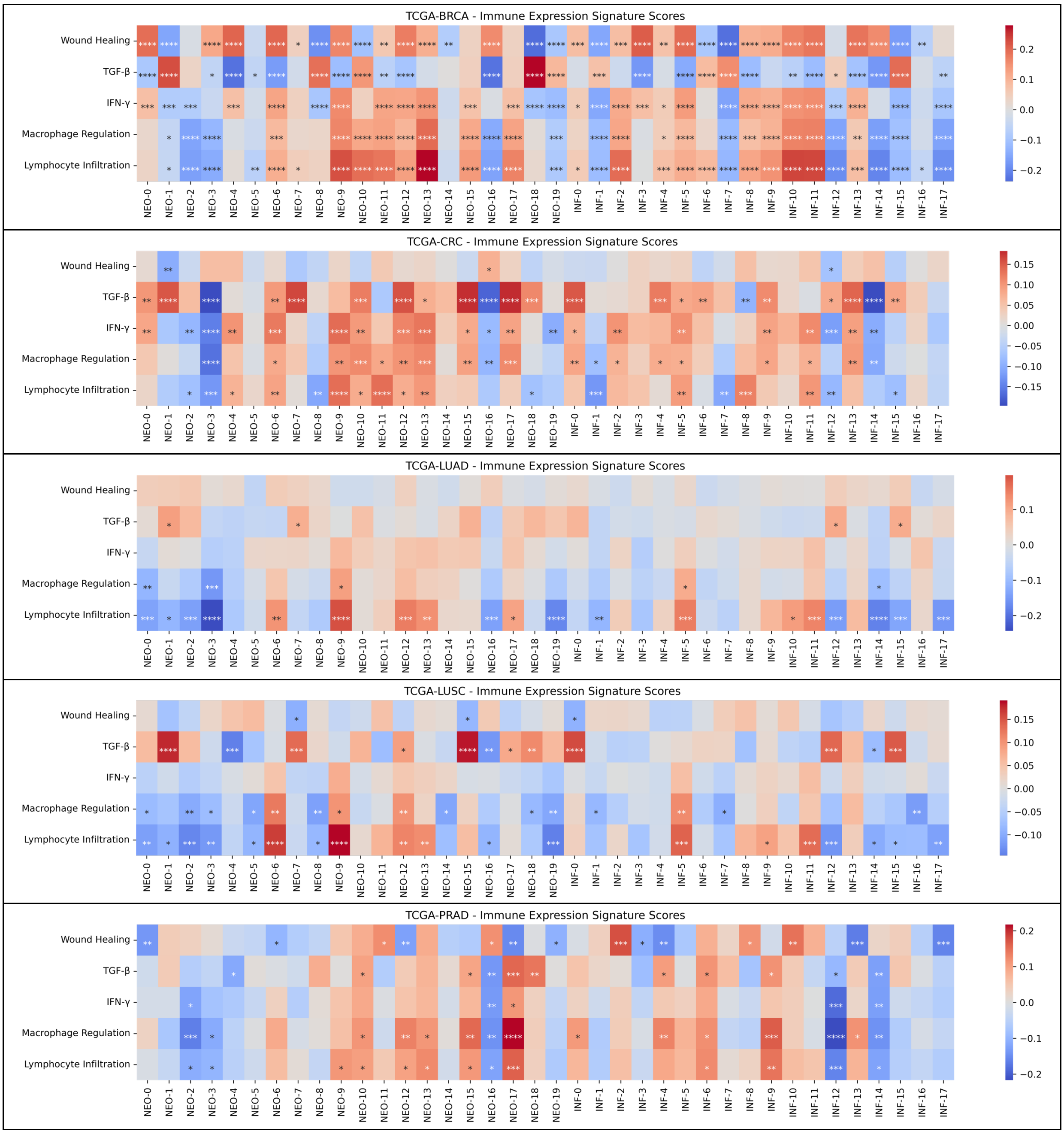


**S7:** Associations between immune expression signature scores and Star-Motifs – Kendall's tau correlation coefficients, FDR BH correction. *P≤0.05, **P≤0.01, ***P≤0.001, ****P≤0.0001.

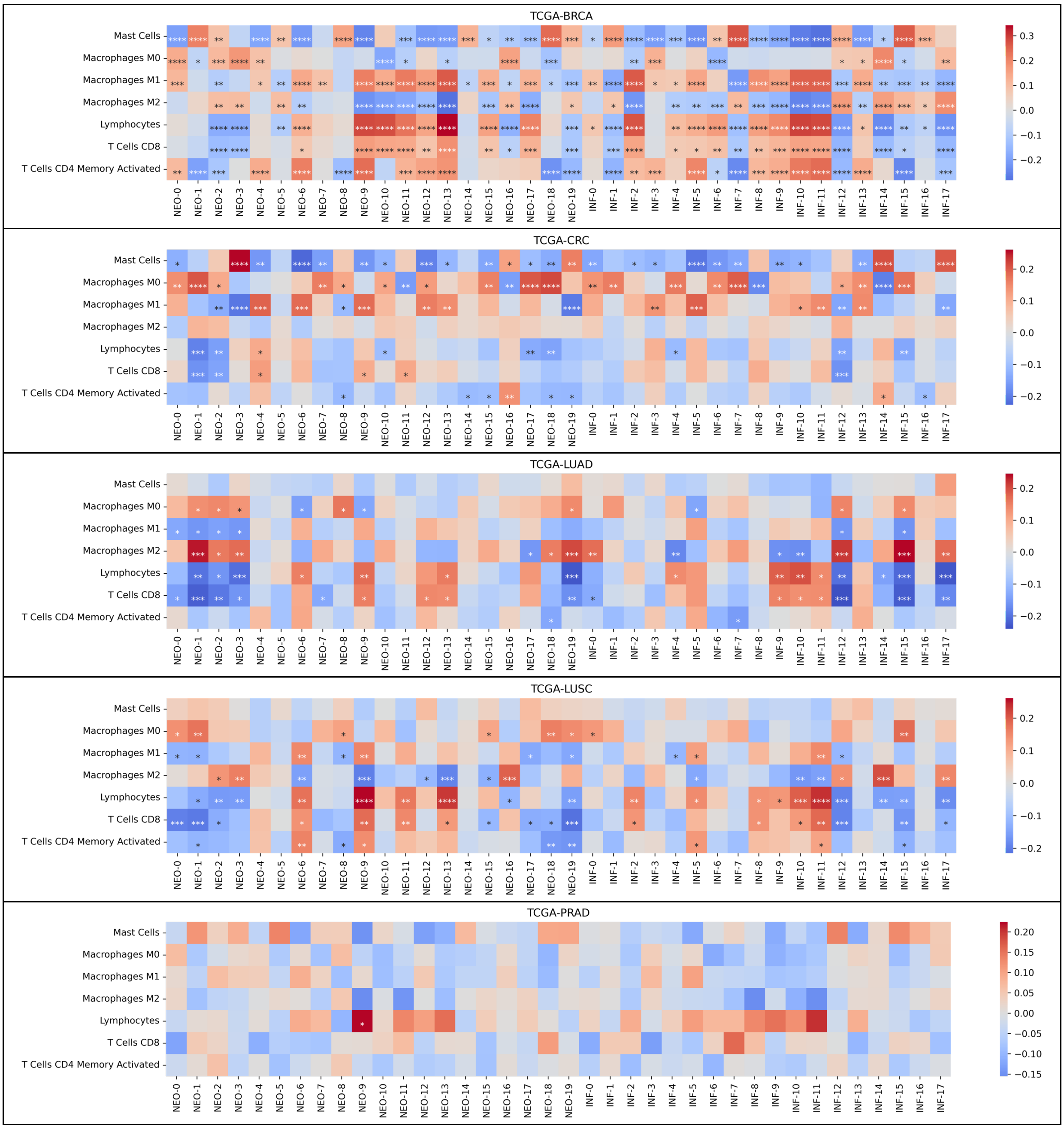


**S8:** Associations between molecular estimates of immune cells and Star-Motifs – Kendall's tau correlation coefficients, FDR BH correction. *P≤0.05, **P≤0.01, ***P≤0.001, ****P≤0.0001.

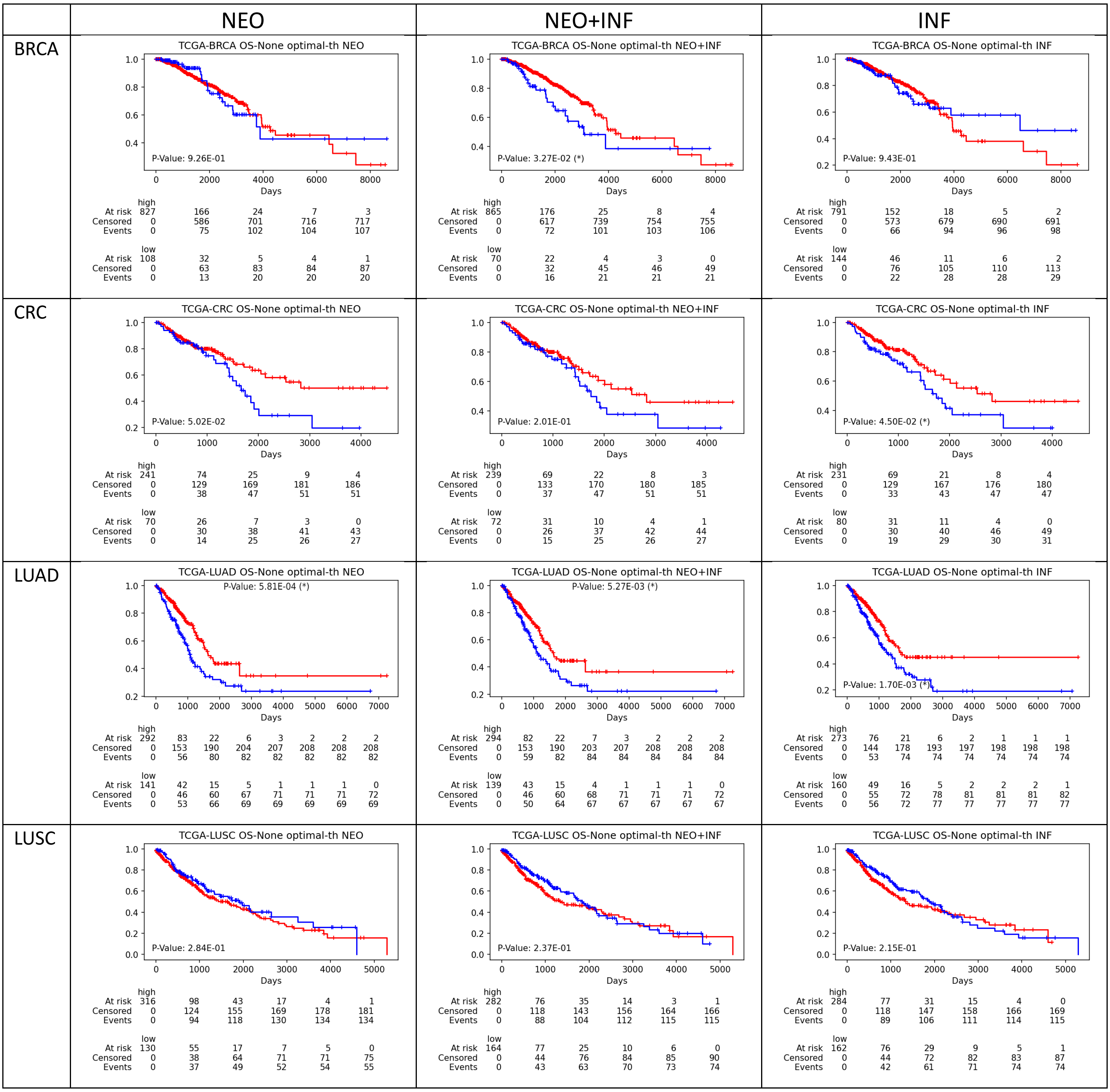


**S9:** Kaplan-Meier analysis of stratification of low (blue) and high (red) survival patients via RSF model outputs on test sets. Plots show the experiments with different input combinations (NEO Star-Motif frequencies, INF Star-Motif frequencies, NEO+INF Star-Motif frequencies) across cancer types. Low and high survival are defined by optimal thresholds selected on model predictions on the discovery sets at each fold. Log-rank test was used to test for statistical significance in survival distributions between patient groups (*p-value < 0.05).

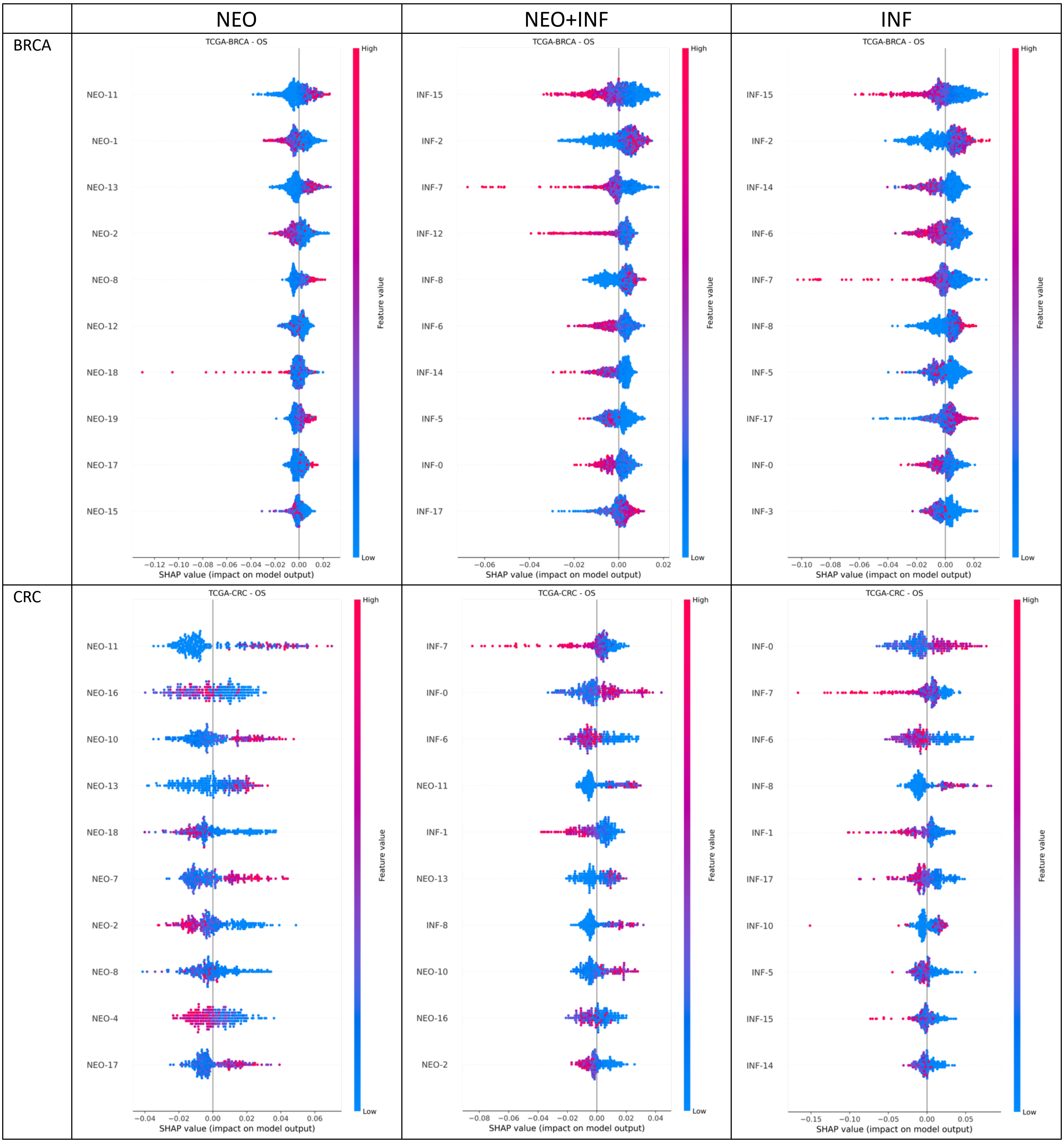


**S10:** SHAP (SHapley Additive exPlanations) plots for OS prediction on BRCA and CRC with different motif sets as input. Plots visualise the contributions of Star-Motifs in the model predictions, showing the top ten contributing the most. A positive SHAP value (on the right side) is associated with better survival, while colours indicate the feature value, high (red) or low (blue), associated with the SHAP scores.

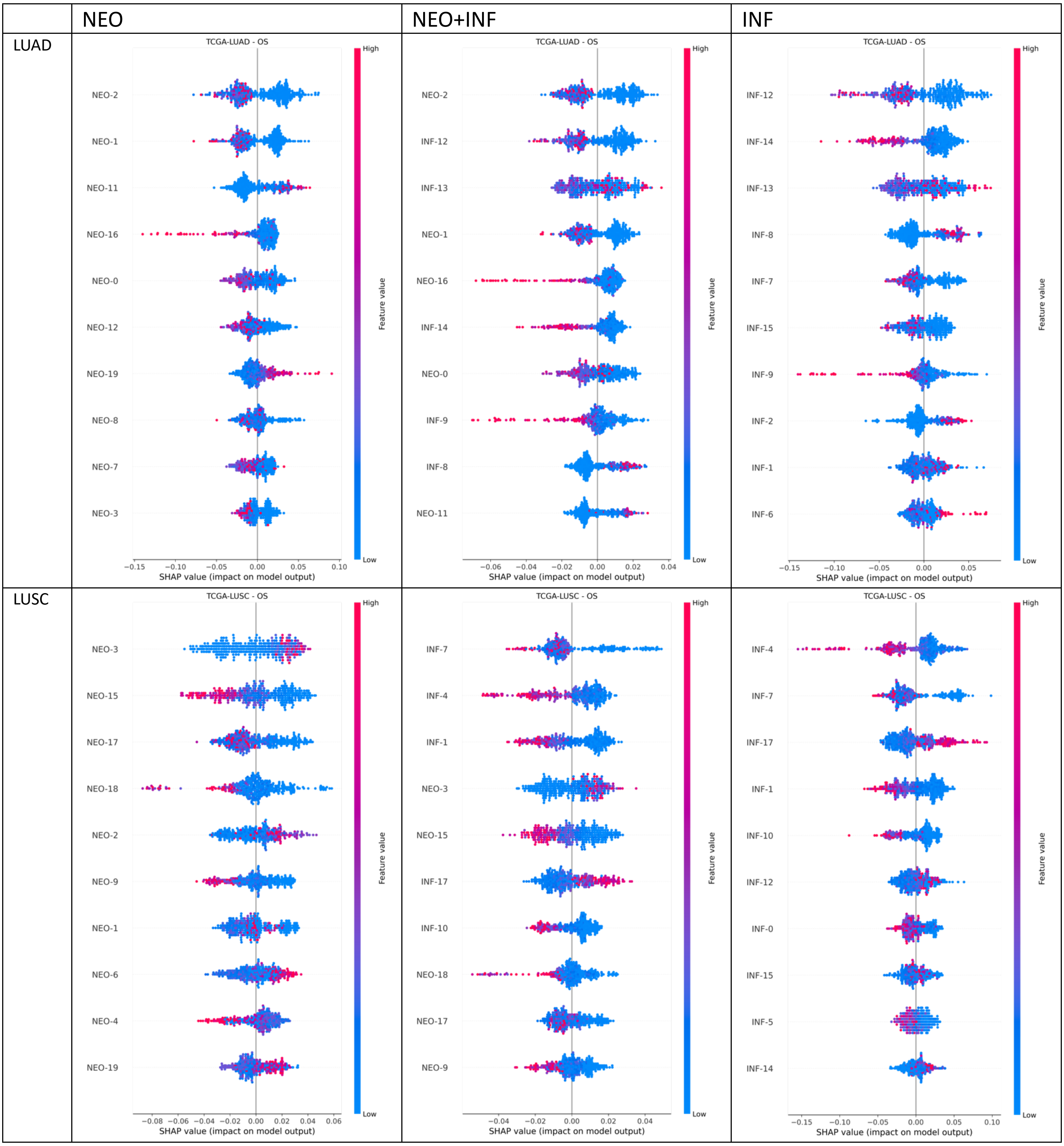


**S11:** SHAP (SHapley Additive exPlanations) plots for OS prediction on LUAD and LUSC with different motif sets as input. Plots visualise the contributions of Star-Motifs in the model predictions, showing the top ten contributing the most. A positive SHAP value (on the right side) is associated with better survival, while colours indicate the feature value, high (red) or low (blue), associated with the SHAP scores.

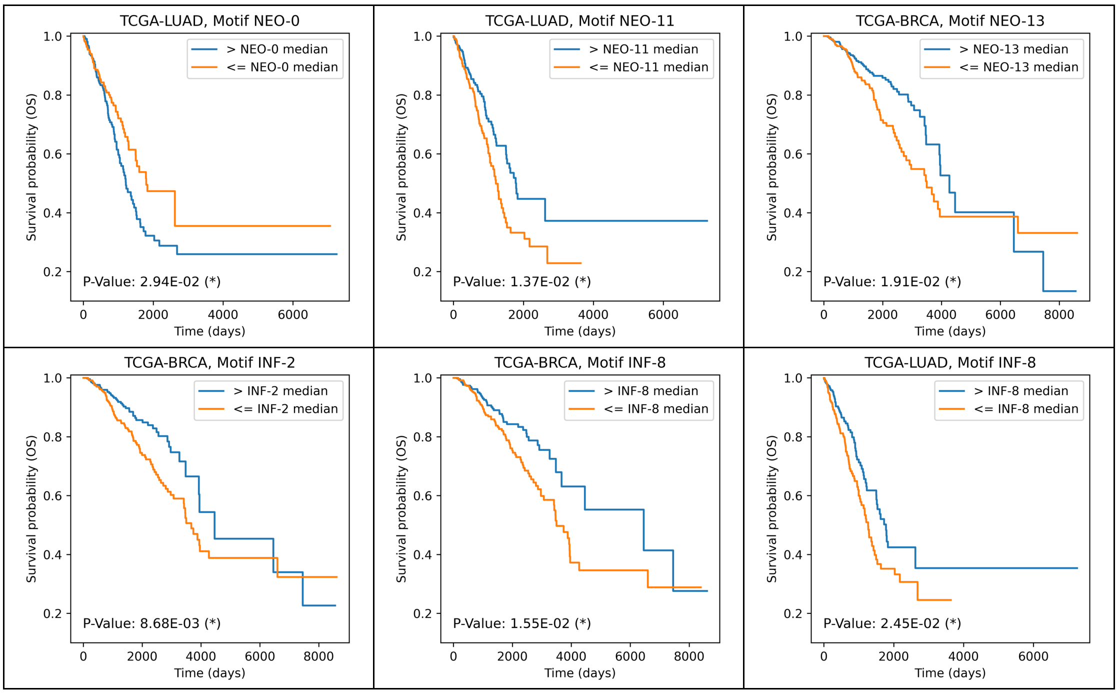


**S12:** Plots showing Kaplan–Meier curves for OS according to patient stratifications using the median values of individual Star-Motif frequencies as the threshold, together with corresponding bootstrap corrected log-rank p-values.

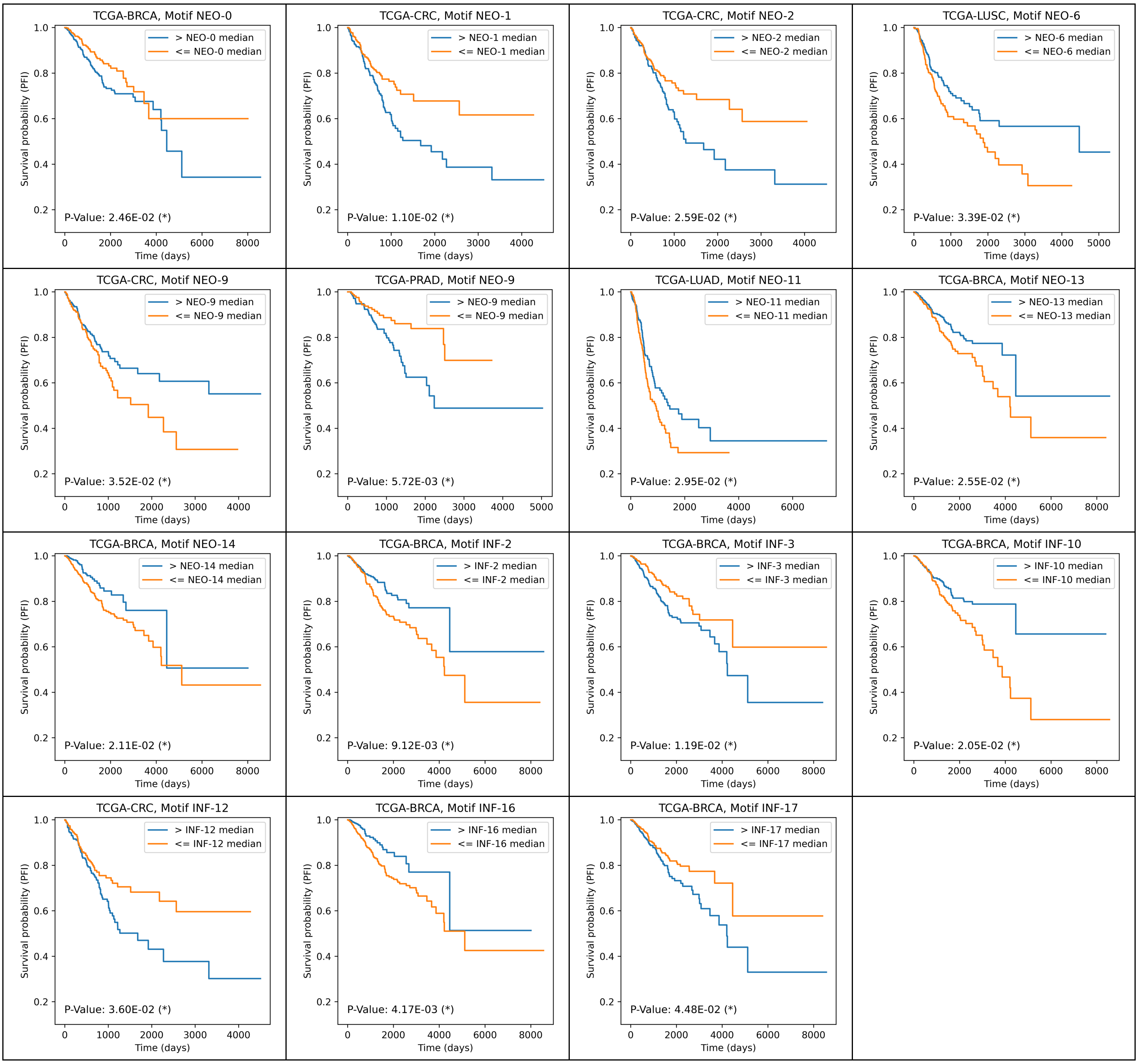


**S13:** Plots showing Kaplan–Meier curves for PFI according to patient stratifications using the median values of individual Star-Motif frequencies as the threshold, together with corresponding bootstrap corrected log-rank p-values.

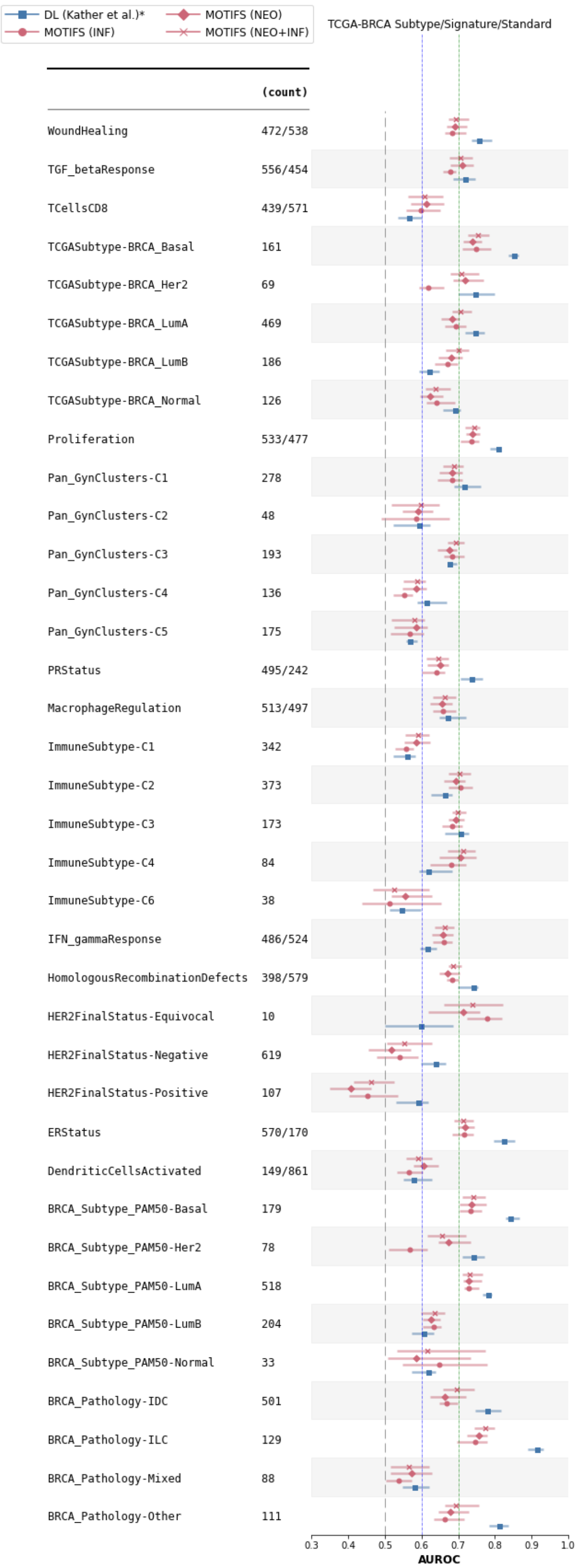


| | (count) | KATHER | NEO | INF | NEO+INF |
|---|---|---|---|---|---|
| WoundHealing | 472/538 | 0.755 | 0.691 | 0.684 | 0.693 |
| TGF_betaResponse | 556/454 | 0.719 | 0.71 | 0.678 | 0.707 |
| TCellsCD8 | 439/571 | 0.565 | 0.612 | 0.599 | 0.608 |
| TCGASubtype-BRCA_Basal | 161 | 0.852 | 0.738 | 0.748 | 0.753 |
| TCGASubtype-BRCA_Her2 | 69 | 0.745 | 0.719 | 0.618 | 0.709 |
| TCGASubtype-BRCA_LumA | 469 | 0.747 | 0.682 | 0.694 | 0.707 |
| TCGASubtype-BRCA_LumB | 186 | 0.621 | 0.681 | 0.672 | 0.7 |
| TCGASubtype-BRCA_Normal | 126 | 0.692 | 0.622 | 0.641 | 0.639 |
| Proliferation | 533/477 | 0.81 | 0.739 | 0.736 | 0.743 |
| Pan_GynClusters-C1 | 278 | 0.717 | 0.683 | 0.682 | 0.688 |
| Pan_GynClusters-C2 | 48 | 0.592 | 0.59 | 0.585 | 0.597 |
| Pan_GynClusters-C3 | 193 | 0.676 | 0.677 | 0.683 | 0.692 |
| Pan_GynClusters-C4 | 136 | 0.612 | 0.585 | 0.553 | 0.587 |
| Pan_GynClusters-C5 | 175 | 0.569 | 0.585 | 0.568 | 0.581 |
| PRStatus | 495/242 | 0.737 | 0.651 | 0.64 | 0.645 |
| MacrophageRegulation | 513/497 | 0.671 | 0.657 | 0.657 | 0.663 |
| ImmuneSubtype-C1 | 342 | 0.56 | 0.584 | 0.557 | 0.59 |
| ImmuneSubtype-C2 | 373 | 0.664 | 0.692 | 0.706 | 0.704 |
| ImmuneSubtype-C3 | 173 | 0.706 | 0.693 | 0.684 | 0.699 |
| ImmuneSubtype-C4 | 84 | 0.618 | 0.707 | 0.681 | 0.714 |
| ImmuneSubtype-C6 | 38 | 0.545 | 0.556 | 0.512 | 0.524 |
| IFN_gammaResponse | 486/524 | 0.616 | 0.658 | 0.661 | 0.664 |
| HomologousRecombinationDefects | 398/579 | 0.742 | 0.672 | 0.682 | 0.687 |
| HER2FinalStatus-Equivocal | 10 | 0.598 | 0.714 | 0.779 | 0.739 |
| HER2FinalStatus-Negative | 619 | 0.638 | 0.518 | 0.541 | 0.552 |
| HER2FinalStatus-Positive | 107 | 0.591 | 0.408 | 0.452 | 0.462 |
| ERStatus | 570/170 | 0.824 | 0.718 | 0.716 | 0.712 |
| DendriticCellsActivated | 149/861 | 0.578 | 0.605 | 0.566 | 0.591 |
| BRCA_Subtype_PAM50-Basal | 179 | 0.841 | 0.737 | 0.733 | 0.74 |
| BRCA_Subtype_PAM50-Her2 | 78 | 0.74 | 0.674 | 0.568 | 0.656 |
| BRCA_Subtype_PAM50-LumA | 518 | 0.781 | 0.73 | 0.728 | 0.731 |
| BRCA_Subtype_PAM50-LumB | 204 | 0.606 | 0.626 | 0.632 | 0.637 |
| BRCA_Subtype_PAM50-Normal | 33 | 0.619 | 0.585 | 0.649 | 0.617 |
| BRCA_Pathology-IDC | 501 | 0.779 | 0.664 | 0.667 | 0.695 |
| BRCA_Pathology-ILC | 129 | 0.915 | 0.755 | 0.745 | 0.773 |
| BRCA_Pathology-Mixed | 88 | 0.581 | 0.573 | 0.539 | 0.566 |
| BRCA_Pathology-Other | 111 | 0.812 | 0.679 | 0.664 | 0.694 |

**S14:** Comprehensive results of the standard/signature/subtype prediction experiments in the **TCGA-BRCA**, illustrating how random forest models trained on Star-Motif frequency vectors (NEO, INF, or both combined) perform relative to the DL benchmark.

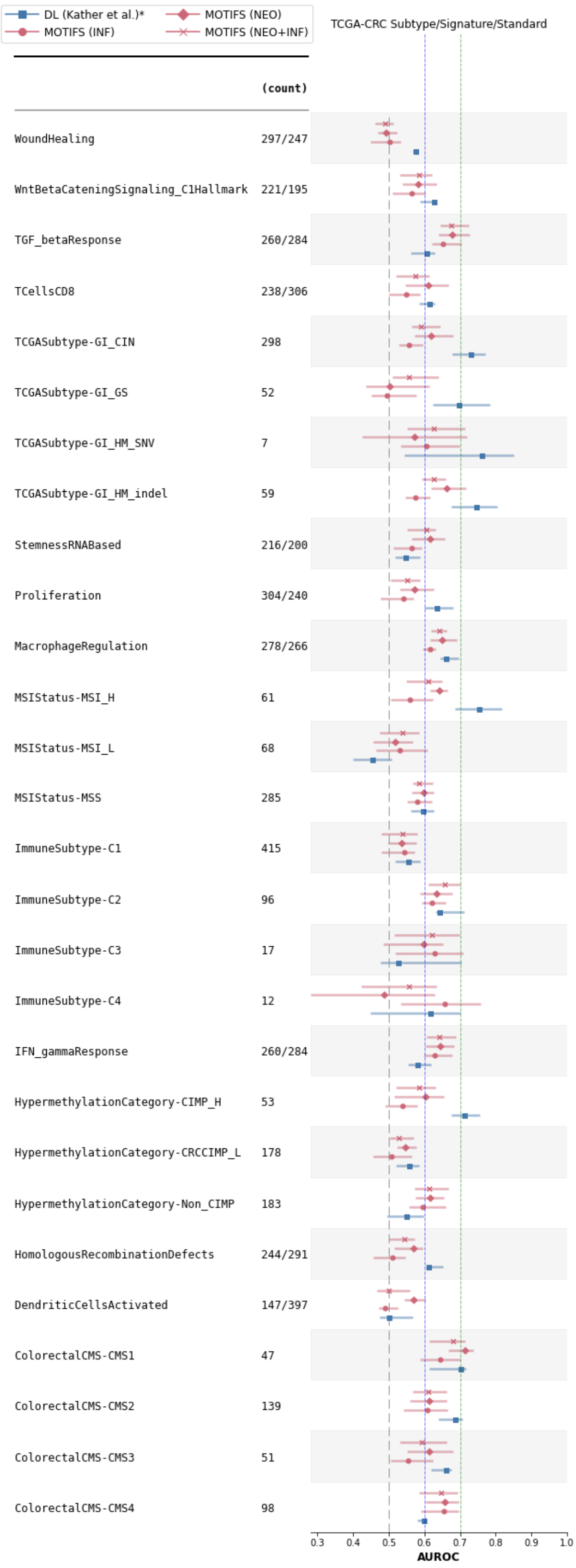


| | (count) | KATHER | NEO | INF | NEO+INF |
|---|---|---|---|---|---|
| WoundHealing | 297/247 | 0.575 | 0.492 | 0.503 | 0.489 |
| WntBetaCateningSignaling_C1Hallmark | 221/195 | 0.626 | 0.582 | 0.565 | 0.585 |
| TGF_betaResponse | 260/284 | 0.607 | 0.678 | 0.652 | 0.676 |
| TCellsCD8 | 238/306 | 0.614 | 0.611 | 0.549 | 0.574 |
| TCGASubtype-GI_CIN | 298 | 0.73 | 0.619 | 0.556 | 0.59 |
| TCGASubtype-GI_GS | 52 | 0.697 | 0.503 | 0.496 | 0.557 |
| TCGASubtype-GI_HM_SNV | 7 | 0.761 | 0.572 | 0.606 | 0.627 |
| TCGASubtype-GI_HM_indel | 59 | 0.745 | 0.662 | 0.574 | 0.626 |
| StemnessRNABased | 216/200 | 0.546 | 0.617 | 0.564 | 0.605 |
| Proliferation | 304/240 | 0.635 | 0.571 | 0.542 | 0.551 |
| MacrophageRegulation | 278/266 | 0.659 | 0.651 | 0.617 | 0.641 |
| MSIStatus-MSI_H | 61 | 0.752 | 0.642 | 0.56 | 0.611 |
| MSIStatus-MSI_L | 68 | 0.455 | 0.519 | 0.531 | 0.539 |
| MSIStatus-MSS | 285 | 0.596 | 0.597 | 0.58 | 0.584 |
| ImmuneSubtype-C1 | 415 | 0.554 | 0.536 | 0.544 | 0.538 |
| ImmuneSubtype-C2 | 96 | 0.642 | 0.634 | 0.62 | 0.658 |
| ImmuneSubtype-C3 | 17 | 0.527 | 0.597 | 0.628 | 0.622 |
| ImmuneSubtype-C4 | 12 | 0.615 | 0.488 | 0.658 | 0.556 |
| IFN_gammaResponse | 260/284 | 0.58 | 0.644 | 0.628 | 0.642 |
| HypermethylationCategory-CIMP_H | 53 | 0.712 | 0.602 | 0.538 | 0.586 |
| HypermethylationCategory-CRCCIMP_L | 178 | 0.558 | 0.547 | 0.507 | 0.528 |
| HypermethylationCategory-Non_CIMP | 183 | 0.548 | 0.615 | 0.595 | 0.613 |
| HomologousRecombinationDefects | 244/291 | 0.611 | 0.569 | 0.511 | 0.543 |
| DendriticCellsActivated | 147/397 | 0.5 | 0.571 | 0.491 | 0.501 |
| ColorectalCMS-CMS1 | 47 | 0.7 | 0.714 | 0.645 | 0.68 |
| ColorectalCMS-CMS2 | 139 | 0.686 | 0.613 | 0.608 | 0.611 |
| ColorectalCMS-CMS3 | 51 | 0.659 | 0.613 | 0.554 | 0.592 |
| ColorectalCMS-CMS4 | 98 | 0.598 | 0.658 | 0.655 | 0.647 |

**S15:** Comprehensive results of the standard/signature/subtype prediction experiments in the **TCGA-CRC**, illustrating how random forest models trained on Star-Motif frequency vectors (NEO, INF, or both combined) perform relative to the DL benchmark.

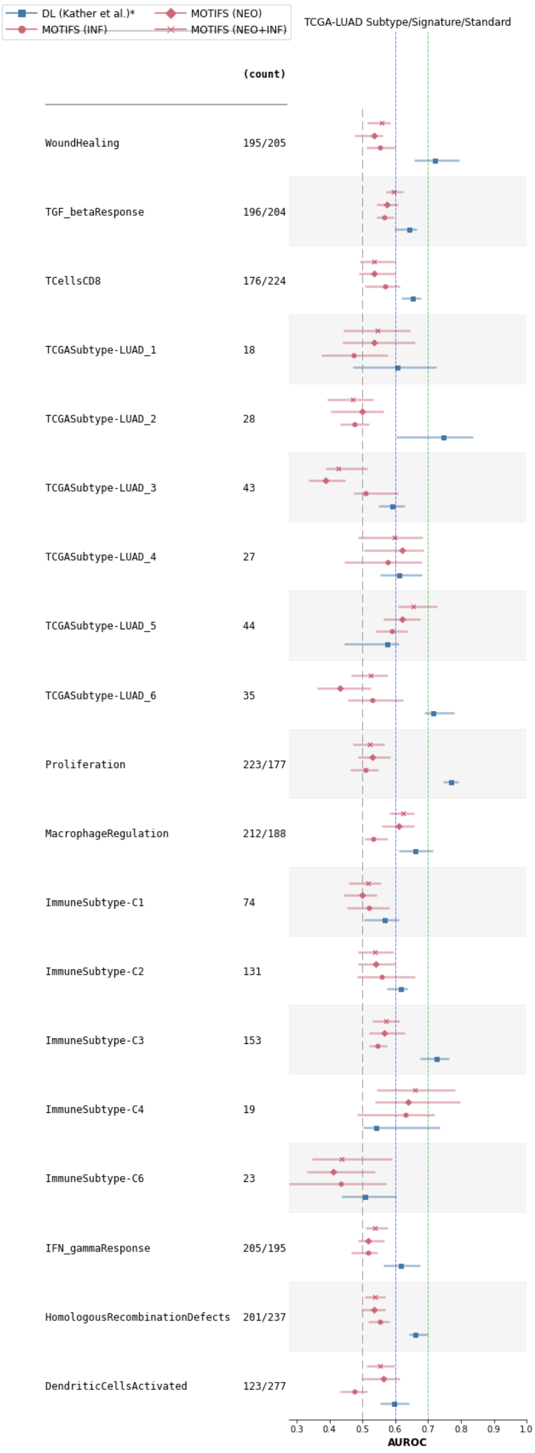


| | (count) | KATHER | NEO | INF | NEO+INF |
|---|---|---|---|---|---|
| WoundHealing | 195/205 | 0.719 | 0.535 | 0.554 | 0.559 |
| TGF_betaResponse | 196/204 | 0.643 | 0.574 | 0.568 | 0.596 |
| TCellsCD8 | 176/224 | 0.653 | 0.536 | 0.57 | 0.535 |
| TCGASubtype-LUAD_1 | 18 | 0.606 | 0.535 | 0.475 | 0.547 |
| TCGASubtype-LUAD_2 | 28 | 0.746 | 0.5 | 0.475 | 0.471 |
| TCGASubtype-LUAD_3 | 43 | 0.589 | 0.388 | 0.51 | 0.427 |
| TCGASubtype-LUAD_4 | 27 | 0.612 | 0.621 | 0.576 | 0.598 |
| TCGASubtype-LUAD_5 | 44 | 0.575 | 0.621 | 0.59 | 0.654 |
| TCGASubtype-LUAD_6 | 35 | 0.715 | 0.432 | 0.532 | 0.525 |
| Proliferation | 223/177 | 0.77 | 0.531 | 0.51 | 0.523 |
| MacrophageRegulation | 212/188 | 0.66 | 0.612 | 0.534 | 0.624 |
| ImmuneSubtype-C1 | 74 | 0.567 | 0.5 | 0.521 | 0.519 |
| ImmuneSubtype-C2 | 131 | 0.617 | 0.541 | 0.558 | 0.538 |
| ImmuneSubtype-C3 | 153 | 0.725 | 0.568 | 0.545 | 0.573 |
| ImmuneSubtype-C4 | 19 | 0.542 | 0.64 | 0.631 | 0.66 |
| ImmuneSubtype-C6 | 23 | 0.507 | 0.411 | 0.435 | 0.437 |
| IFN_gammaResponse | 205/195 | 0.617 | 0.517 | 0.518 | 0.538 |
| HomologousRecombinationDefects | 201/237 | 0.66 | 0.535 | 0.553 | 0.539 |
| DendriticCellsActivated | 123/277 | 0.596 | 0.564 | 0.477 | 0.553 |

**S16:** Comprehensive results of the standard/signature/subtype prediction experiments in the **TCGA-LUAD**, illustrating how random forest models trained on Star-Motif frequency vectors (NEO, INF, or both combined) perform relative to the DL benchmark.

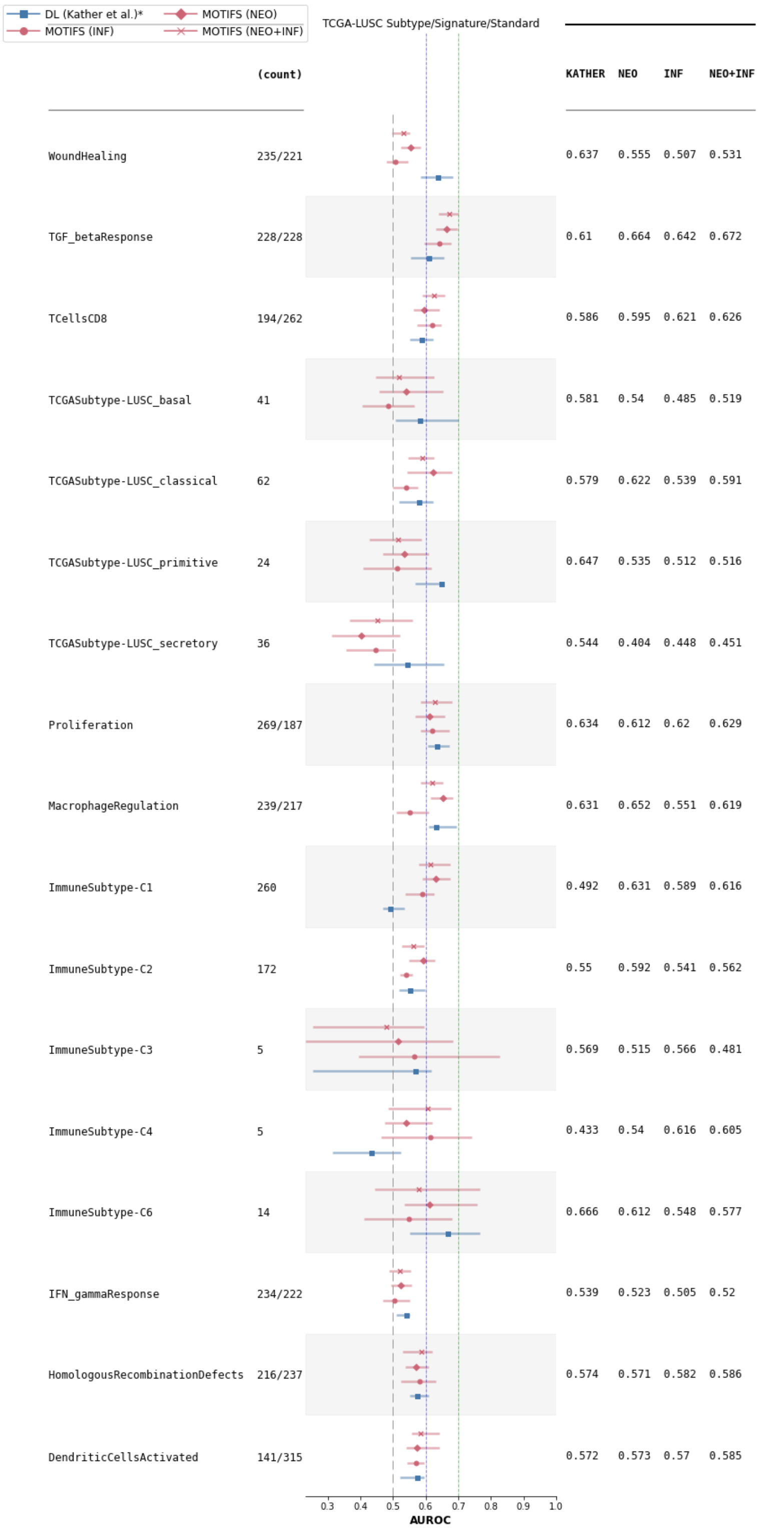


**S17:** Comprehensive results of the standard/signature/subtype prediction experiments in the **TCGA-LUSC**, illustrating how random forest models trained on Star-Motif frequency vectors (NEO, INF, or both combined) perform relative to the DL benchmark.

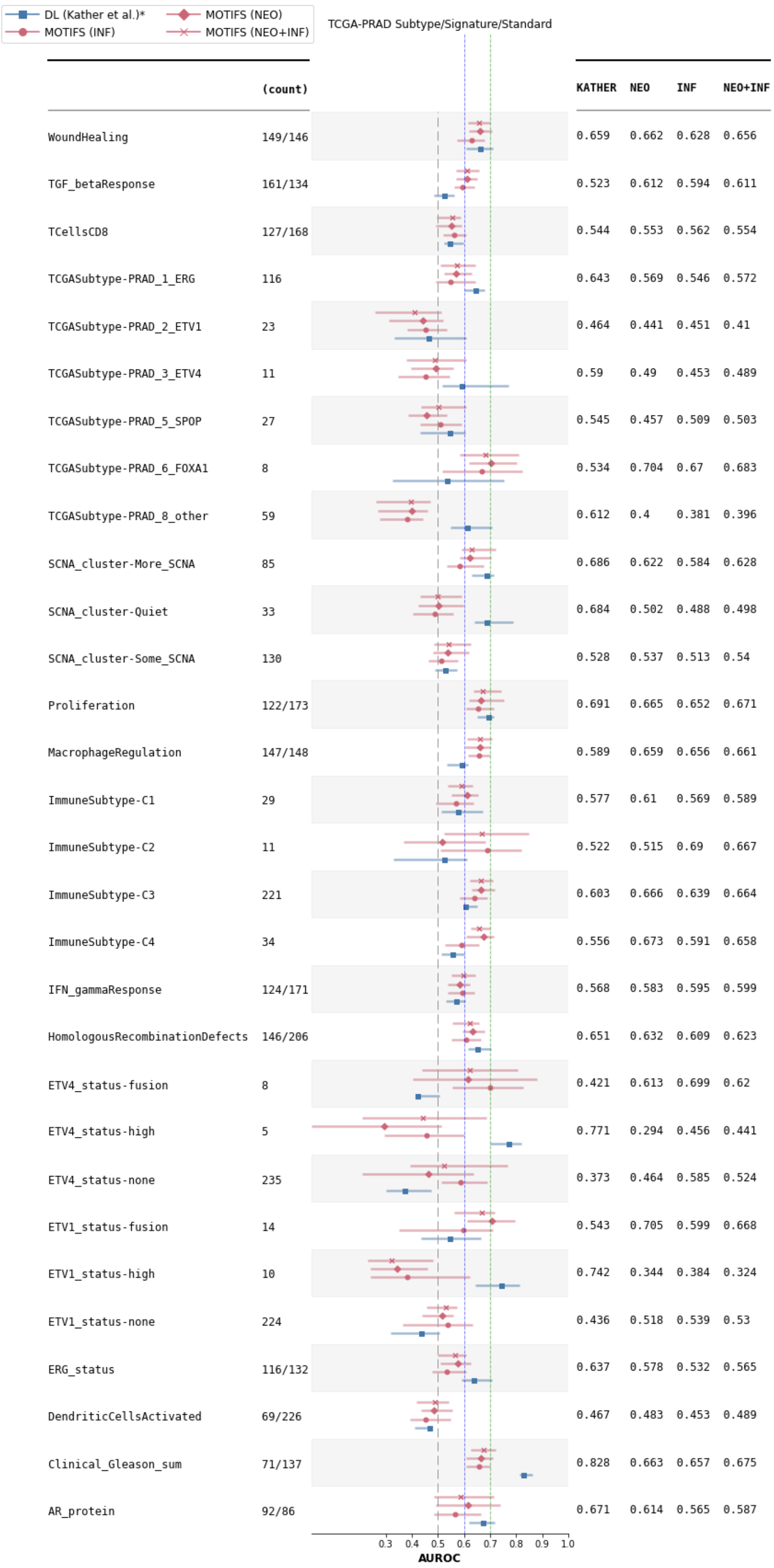


| | (count) | KATHER | NEO | INF | NEO+INF |
|---|---|---|---|---|---|
| WoundHealing | 149/146 | 0.659 | 0.662 | 0.628 | 0.656 |
| TGF_betaResponse | 161/134 | 0.523 | 0.612 | 0.594 | 0.611 |
| TCellsCD8 | 127/168 | 0.544 | 0.553 | 0.562 | 0.554 |
| TCGASubtype-PRAD_1_ERG | 116 | 0.643 | 0.569 | 0.546 | 0.572 |
| TCGASubtype-PRAD_2_ETV1 | 23 | 0.464 | 0.441 | 0.451 | 0.41 |
| TCGASubtype-PRAD_3_ETV4 | 11 | 0.59 | 0.49 | 0.453 | 0.489 |
| TCGASubtype-PRAD_5_SPOP | 27 | 0.545 | 0.457 | 0.509 | 0.503 |
| TCGASubtype-PRAD_6_FOXA1 | 8 | 0.534 | 0.704 | 0.67 | 0.683 |
| TCGASubtype-PRAD_8_other | 59 | 0.612 | 0.4 | 0.381 | 0.396 |
| SCNA_cluster-More_SCNA | 85 | 0.686 | 0.622 | 0.584 | 0.628 |
| SCNA_cluster-Quiet | 33 | 0.684 | 0.502 | 0.488 | 0.498 |
| SCNA_cluster-Some_SCNA | 130 | 0.528 | 0.537 | 0.513 | 0.54 |
| Proliferation | 122/173 | 0.691 | 0.665 | 0.652 | 0.671 |
| MacrophageRegulation | 147/148 | 0.589 | 0.659 | 0.656 | 0.661 |
| ImmuneSubtype-C1 | 29 | 0.577 | 0.61 | 0.569 | 0.589 |
| ImmuneSubtype-C2 | 11 | 0.522 | 0.515 | 0.69 | 0.667 |
| ImmuneSubtype-C3 | 221 | 0.603 | 0.666 | 0.639 | 0.664 |
| ImmuneSubtype-C4 | 34 | 0.556 | 0.673 | 0.591 | 0.658 |
| IFN_gammaResponse | 124/171 | 0.568 | 0.583 | 0.595 | 0.599 |
| HomologousRecombinationDefects | 146/206 | 0.651 | 0.632 | 0.609 | 0.623 |
| ETV4_status-fusion | 8 | 0.421 | 0.613 | 0.699 | 0.62 |
| ETV4_status-high | 5 | 0.771 | 0.294 | 0.456 | 0.441 |
| ETV4_status-none | 235 | 0.373 | 0.464 | 0.585 | 0.524 |
| ETV1_status-fusion | 14 | 0.543 | 0.705 | 0.599 | 0.668 |
| ETV1_status-high | 10 | 0.742 | 0.344 | 0.384 | 0.324 |
| ETV1_status-none | 224 | 0.436 | 0.518 | 0.539 | 0.53 |
| ERG_status | 116/132 | 0.637 | 0.578 | 0.532 | 0.565 |
| DendriticCellsActivated | 69/226 | 0.467 | 0.483 | 0.453 | 0.489 |
| Clinical_Gleason_sum | 71/137 | 0.828 | 0.663 | 0.657 | 0.675 |
| AR_protein | 92/86 | 0.671 | 0.614 | 0.565 | 0.587 |

**S18:** Comprehensive results of the standard/signature/subtype prediction experiments in the **TCGA-PRAD**, illustrating how random forest models trained on Star-Motif frequency vectors (NEO, INF, or both combined) perform relative to the DL benchmark.

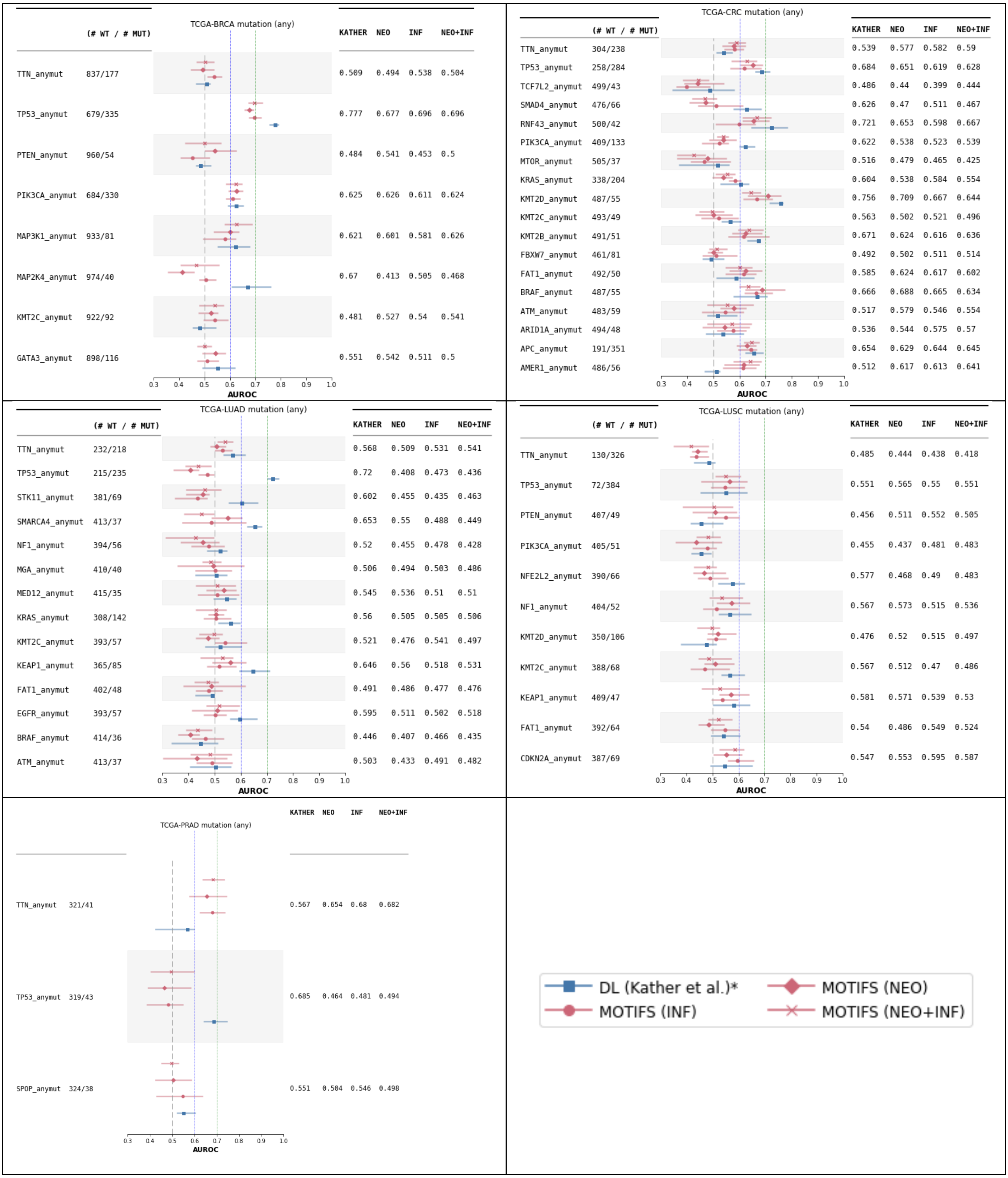


**S19:** Mutation prediction results for each of the five cancer cohorts (BRCA, CRC, LUAD, LUSC, PRAD), showing how random forest models trained on Star-Motif frequency vectors (NEO, INF, or both combined) perform relative to the DL benchmark. Although not all mutations are predictable, several (e.g., *MAP3K1 and PIK3CA* on BRCA; *FAT1, BRAF, APC, and AMER1* on CRC; *TTN* on PRAD) show comparable or higher performance with Star-Motifs.

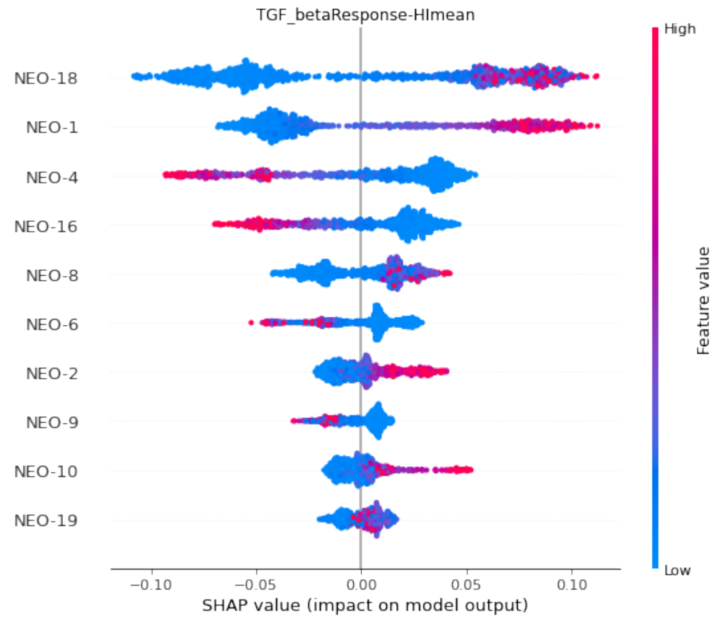


**S20:** The figure shows the SHAP (SHapley Additive exPlanations) plot for the binary prediction of TGF-β (high, low) on TCGA-BRCA, which can be used to identify Star-Motifs that contribute most to the prediction. A positive SHAP value (on the right side) is associated with high TGF-β, while colours indicate the feature value, high (red) or low (blue), associated with the SHAP scores.

NEO

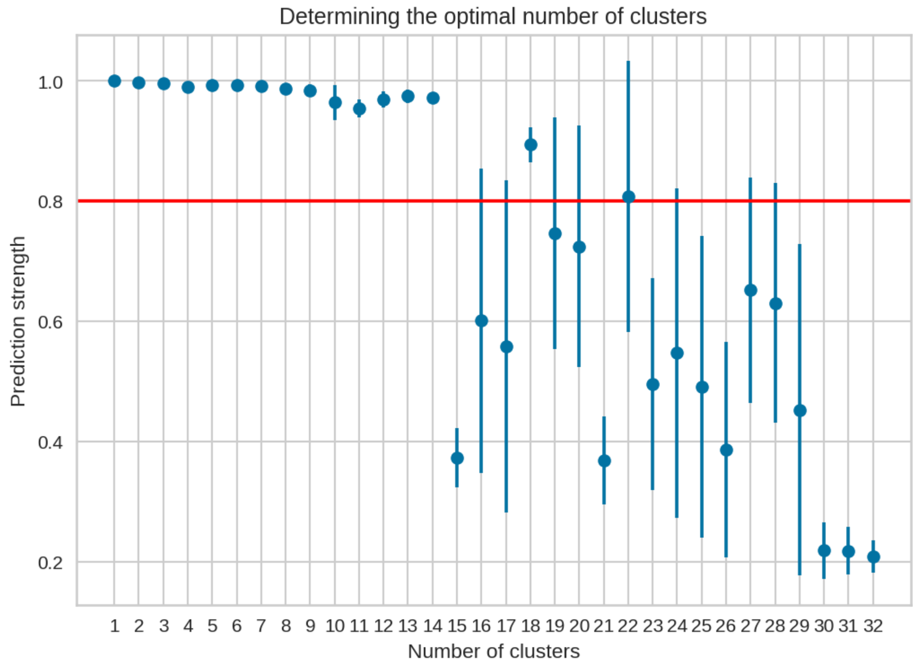


INF

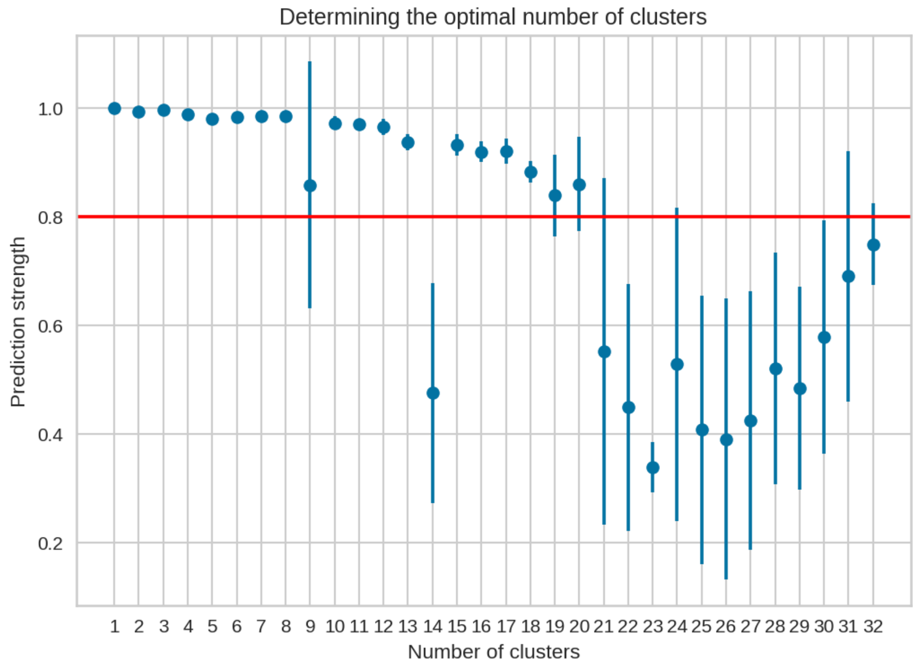


**S21:** Plots (NEO and INF separately) showing prediction strength values (mean and std over 5 runs) for each tested number of clusters (k) value. Using this approach, the optimal number of clusters (maximum k with mean prediction strength > 0.8) is selected as 20 for INF and 22 for NEO.
